# The eIF4E-homologous protein 4EHP (nCBP) regulates thermotolerance by modulating heat-responsive mRNAs and the HSP repertoire in Arabidopsis thaliana

**DOI:** 10.64898/2026.04.09.717490

**Authors:** Marlon A. Pulido-Torres, Karen Quiroz-Núñez, Eduardo M. Dorado-Cruz, Susana De la Torre-Díaz, Abraham León-Domínguez, Jorge Herrera-Díaz, Oliver Herzog, Héctor Quezada, Manuela Nájera-Martínez, Magdalena Weingartner, Tzvetanka D. Dinkova

## Abstract

Translation initiation factors of the eIF4E family play a crucial role in regulating translation and the cellular metabolism of mRNAs. Research has addressed the role of canonical 4E family isoforms in development, stress response, and during viral infection. Nevertheless, the class-II eIF4E family member cap-binding protein 4EHP (nCBP), has remained poorly characterized in plant stress responses.

In this study, we show that loss of 4EHP confers enhanced basal and acquired thermotolerance and causes a mild flowering delay without major root defects. Under heat stress, 4EHP-GFP re-localizes from a diffuse cytosolic pattern to cytoplasmatic foci, co-localizing with canonical stress granule (SG) markers. Transcriptomic analysis under control, acclimation and heat stress conditions reveals that 4EHP limits the accumulation of heat-responsive mRNAs, especially those encoding heat shock proteins (HSPs), which remain constitutively expressed in *4ehp-1* mutant under control and heat stress conditions. Proteomic analysis also indicates that the absence of 4EHP alters the repertoire of HSPs compared with wild type (Col-0), especially upon heat stress, without significantly impairing the recruitment of corresponding mRNAs to translationally active polysomes. Together, our results indicate that 4EHP negatively modulates the accumulation of a specific subset of heat-responsive mRNAs fine-tuning chaperone production, via heat-responsive SG regulatory pathways.

## Introduction

Due to their sessile nature, plants are unable to avoid environmental stress. Hence, they have developed a set of sophisticated mechanisms to respond appropriately to ever changing surrounding conditions in place (reviewed in Ding *et al*., 2020). An important step for their adaptation is the control of gene expression, allowing appropriate response, metabolism, growth rate, and development readjustment to withstand stress and enhance survivability (Rao, 2006). Albeit transcription readjustment is required to contend with stress; multiple levels of expression regulation operate to exert required responses. Regulation of translation is especially important since the process is high energy-demanding and a key step for modulating protein levels (Schwanhausser *et al*., 2011).

The eukaryotic initiation factor 4E (eIF4E) protein family activates mRNAs during translation initiation. These “L” shaped proteins recognize the m7GpppN (5′cap) in mature mRNAs through conserved aromatic residues (Matsuo *et al*., 1997). eIF4E interacts with initiation factors eIF4G and eIF4A forming the heterotrimeric complex eIF4F. Parallel to eIF4F formation on mRNA recognition, the 43S pre-initiation complex (PIC) assembly takes place, through association of 40S ribosomal subunit, eIF1, eIF1A, eIF3 and eIF5 with the ternary complex eIF2/GTP/Met-tRNA_i_^Met^. Activated mRNAs are joined by PIC through interaction between eIF4G and eIF3, then become circularized by eIF4G-PABP (poly A binding protein) association to form the 48S initiation complex, which scans towards the AUG start codon (Preiss & M, 2003).

eIF4E proteins have been grouped in Class I and II, depending on the conservation of Trp residues at positions 43 and 56 according to human eIF4E (Joshi *et al*., 2005). *Arabidopsis thaliana* Class I comprises eIF4E1, eIF4E1b, eIF4E1c and eIFiso4E. eIF4E1 and eIFiso4E form complexes with eIF4G and eIFiso4G, respectively, and support general translation (Browning & Bailey-Serres, 2015). They display partially redundant functions but may contribute to selective translation of mRNAs under stress (Browning & Bailey-Serres, 2015). Class II includes nCBP, for novel Cap-binding protein (Ruud *et al*., 1998), conserved in fungi and animals (Joshi *et al*., 2005), and widely referred as 4E homologous protein 4EHP (Christie & Igreja, 2023). To follow up on this more general designation we will hereafter refer to this protein as 4EHP.

In animals, 4EHP has a role in selective translation of specific mRNAs at certain developmental stages or under hypoxia stress. It has also been identified as part of translation repression complexes, where, in association with diverse partners, it is required to inhibit translation of specific mRNAs (reviewed in Christie & Igreja, 2023). Although 4EHP was first described in plants, little is known about its role in these organisms. *Arabidopsis* 4EHP is expressed at lower levels than Class I eIF4Es, and its protein accumulation is mainly induced in actively growing tissues (Bush *et al*., 2009). Interestingly, 4EHP shows a 4- to 10-fold lower affinity for cap analogs than eIF4E (Kropiwnicka *et al*., 2015) and displays modest ability to stimulate translation in vitro (Ruud *et al*., 1998). Besides, plant 4EHP has been characterized as a recessive resistance gene for capped RNA virus infection (Suzuki *et al*., 2025) and was recently associated with resistance to insect pests (Chen *et al*., 2023).

Heat stress represents a major challenge to plants, inducing a complex and highly coordinated cellular response to maintain protein homeostasis and to ensure survival (Staacke *et al*., 2025). Upon exposure to elevated temperatures, plants activate heat shock transcription factors (mainly HsfA1s), which function as master regulators to trigger a transcriptional cascade leading to heat-responsive gene expression, including upregulation of heat shock encoding proteins (HSPs) (Ohama *et al*., 2016). HSPs function as essential molecular chaperones to prevent misfolded protein aggregation, facilitate correct refolding, and maintain proteome integrity under stress conditions (Kang *et al*., 2022). These chaperones are indispensable to achieve thermotolerance, but their abundance and duration of expression must be tightly controlled, to avoid deleterious effects on plant fitness (Song *et al*., 2009; Sun *et al*., 2016; Chen *et al*., 2018).

In most eukaryotes including plants, elevated temperature leads to a global inhibition of protein synthesis rates to limit accumulation of newly formed, potentially damaged proteins, and prevent proteotoxic stress. At the same time, proteins involved in homeostasis maintenance, such as HSPs, must remain actively expressed as part of the mechanisms of cell survival (Echevarria-Zomeno *et al*., 2013). Thus, *HSP* translation escapes inhibition in part through regulation at mRNA level by specific sequences or modulating the recruitment of regulatory proteins to the transcript (Yanguez *et al*., 2013; Dannfald *et al*., 2025). In addition, recent discoveries reveal the crucial role of crosstalk between translation inhibition, mRNA decay, and resuming translation upon recovery for plant survival (Merret *et al*., 2017).

Stress granules (SGs) are dynamic ribonucleoprotein compartments formed via liquid-liquid phase separation (LLPS) that play a central role in translational control during abiotic stress (Maruri-Lopez *et al*., 2021). These contain stalled translation preinitiation complexes, mRNAs, RNA-binding proteins (RBPs), and initiation factors including eIF4E, eIF4G, and eIF4A, alongside core components such as UBP1, PAB2/4/8, eEF1Bβ, RBP47, and TSN1/2 (Gutierrez-Beltran *et al*., 2015; Kosmacz *et al*., 2019; Lohmann *et al*., 2024). Recently, Tudor Staphylococcal Nuclease (TSN) was identified as a central scaffold protein that organizes a large intrinsically disordered protein network under non stress conditions, preparing SG components for assembly upon heat stress (Gutierrez-Beltran *et al*., 2021). This study reported eIF4E5 (4EHP) among the TSN-interacting proteins, recruited to SGs during heat stress (HS). Independent reports have also connected plant 4EHP to SGs or processing bodies (PBs) through its interaction with Essential for Potexvirus Accumulation 1 (EXA1), a protein with GYF domain involved in multiple protein-protein interactions (Scheer *et al*., 2021; Chen *et al*., 2022). These findings potentially link 4EHP to SG dynamics. However, the function of this eIF4E family member in plant mRNA regulation has not yet been characterized.

Here we addressed the relevance of *Arabidopsis thaliana* 4EHP in heat stress response. We found that the protein re-localized to stress granules upon heat stress and its absence resulted in enhanced thermotolerance compared with wild type (Col-0) plants. Transcriptomic and proteomic analyses of the *4ehp* mutant compared to Col-0 at control, acclimation, and heat stress temperatures, revealed that 4EHP has a role in negative regulation of HSP-encoding mRNAs mostly after heat stress. Upregulation of several HSPs in the mutant was consistent with the thermotolerant phenotype. Therefore, we propose that 4EHP likely functions as a selective repressor whose sequestration into SGs during thermal stress may represent a novel regulatory mechanism to fine tune HSP synthesis and prevent deleterious chaperone overaccumulation.

## Materials and Methods

### Plant materials and growth conditions

*Arabidopsis thaliana* ecotype Columbia-0 (Col-0) was used as the wild-type line. The T-DNA insertion mutant *4ehp-1* (SALK_131503) was obtained from the Arabidopsis Biological Resource Center. The mutant line *eif(iso)4e-1*, was provided by Dr. Christophe Robaglia from CNRS-CEA Université de la Mediterranée, France (Duprat *et al*., 2002). The *eif4e1* mutant line was donated by Dr. Jean Luc Gallois from INRA, Montfavet, France (Bastet *et al*., 2019). The mutant line *hsp101* described by (Hong & Vierling, 2000; McLoughlin *et al*., 2019) was ordered from the Nottingham Arabidopsis Stock Centre. The free GFP reporter line carried the CDS of GFP under the control of a 35S promoter.

For flowering time measurements, plants were grown in soil under long-day (LD; 16 h light/8 h dark) or short-day (SD; 8 h light/16 h dark) photoperiod conditions at 22°C. Total rosette leaf number at bolting was counted, excluding cauline leaves. For root growth analysis, seedlings were grown vertically on MS plates. Primary root length was measured at indicated time points using ImageJ software (Schneider *et al*., 2012). Lateral root primordia and emerged lateral roots were counted under a dissecting microscope. For cell division analysis in the root apical meristem, tissues were stained with propidium iodide, and the number of dividing cells in the cortical cell layer was quantified using confocal microscopy.

### Generation of complementation lines

The full-length *4EHP* coding sequence was amplified from Col-0 cDNA and cloned into the pEarleyGate103 binary vector (Earley *et al*., 2006) to generate C-terminal GFP fusion constructs under the control of the cauliflower mosaic virus (CaMV) 35S promoter (p35S:4EHP-GFP). For native promoter construct, a 1.0-kb genomic region upstream of the *4EHP* start codon was PCR-amplified and cloned together with the *4EHP* genomic sequence into pMDC107 vector (p4EHP:4EHP-GFP). Primers used for cloning are available in **Supporting information Table S1**. Constructs were transformed into *Agrobacterium tumefaciens* strain GV3101 and introduced into *4ehp-1* plants using the floral dip method. Transformants were selected on MS medium containing 25 μg/mL hygromycin, and homozygous T₃ lines showing single-locus insertion were used for phenotypic analyses.

### Thermotolerance assays

#### Acquired thermotolerance

Seven-day old seedlings grown on MS plates were subjected to a priming heat treatment at 38°C for 90 min in a temperature-controlled incubator, followed by 2 hours recovery period at 22°C. Then seedlings were exposed to a heat stress at 45°C for 2 hours and returned to control conditions. Nine days after treatment, seedling recovery was measured by counting the number of expanded green leaves (including cotyledons) per seedling. Damage categories were defined as follows: 0-2 leaves (severe damage), 3-4 leaves (moderate damage), 5-6 or more leaves (low damage). Leaves from control seedlings maintained at 22°C were analyzed in parallel.

#### Basal thermotolerance

Seeds were grown in darkness at 22°C for 3 days, exposed to 45°C for 2 hours and then returned to control temperature. Hypocotyl lengths were measured 3 days after treatment using ImageJ software. Control hypocotyls maintained at 22°C were measured in parallel.

### Protoplast isolation and transient expression

*Arabidopsis* mesophyll protoplasts isolation and transient expression assays were performed following the protocol described by (Yoo *et al*., 2007) with some modifications. Plants were grown under SD conditions. Leaves from 6-week-old plants were cut into small stripes and incubated in enzyme-containing buffer (500 mM sorbitol, 1 mM CaCl_2_, 0.25% macerozym R10, 1% cellulase R10, 10 mM MES-KOH, pH 5.7) for 2 h at 26 °C in the dark shaking at 60 rpm. Protoplasts were subsequently washed twice with MaMG buffer (450 mM sorbitol, 15 mM MgCl_2_, 5 mM MES-KOH, pH 5.7). Plasmids containing eEF1Bβ1–mCherry and PAB8–mCherry were used as SG markers (Lohmann *et al*., 2023), while DCP1–mCherry was used as PB marker (Weber *et al*., 2008). For transformation, 300 µl of protoplast suspension was gently mixed with 10 µg of plasmid DNA and 330 µl of polyethylene glycol (PEG)-Ca buffer (40% PEG 4000, 200 mM mannitol, 100 mM CaCl_2_) and incubated for 30 min in the dark. PEG was removed by washing the protoplasts twice with 3 ml of wash buffer (154 mM NaCl, 125 mM CaCl_2_, 5 mM KCl, 5 mM glucose, 2 mM MES, pH 5.7) followed by overnight incubation in 3 ml of the same buffer. Expression of constructs cloned into estradiol-induced vectors was started by adding 50 µM estradiol during overnight incubation.

### Confocal microscopy and heat stress treatments

For granule localization, transfected protoplasts or 5-day-old intact seedling roots were exposed to heat stress at 40°C for 35 min in a temperature-controlled incubator. Control samples were maintained at 22°C. For recovery experiments, heat-stressed samples were returned to 22°C for 6 h. Protoplasts were placed on glass slides and immediately observed using a Leica TCS SP8 Confocal Platform (Leica Microsystems). GFP fluorescence was excited at 488 nm and detected at 498-514 nm; mCherry fluorescence was excited at 561 nm and detected at 590-630 nm. All protoplast images are shown as 2D maximum intensity projection from *z*-stacks. For granule quantification, the number of cytoplasmic foci per cell were counted using ImageJ software. Co-localization analysis was performed by counting foci showing overlapping GFP and mCherry signals across 10 foci. Fluorescence intensity line profiles were generated using ImageJ software.

### Nucleic acid isolation and analysis

Genomic DNA was extracted from rosette leaves using the CTAB method (Doyle & Doyle, 1987). Homozygous *4ehp-1* plants were identified by PCR using gene-specific primers flanking the T-DNA insertion site (Supplementary Table S1).

Total RNA was extracted using TRIzol reagent (Invitrogen) according to the manufacturer’s instructions. First-strand cDNA synthesis was performed on 2 μg total RNA with SuperScript III Reverse Transcriptase (Invitrogen) and oligo(dT)₁₈ primers. End-point PCR was performed to assess transcript levels of *eIF4E* family members and neighboring genes. Quantitative real-time PCR (RT-qPCR) was conducted with Maxima SYBR Green/ROX qPCR Master Mix (ThermoFisher) in a 7500 Real-time PCR System (Applied Biosystems). Gene-specific primers are available in Supplementary Table S1). Relative expression levels were calculated with 2^-ΔΔCt^ method (Livak & Schmittgen, 2001) and *ACTIN2* (At3g18780) as reference. Three independent biological replicates were performed for each analysis.

### Polyribosome profiling

Polyribosome profiling was performed as described by (Martinez-Silva *et al*., 2012) with some modifications. Briefly, 2 g of 7-day-old seedlings from same tissues and temperature conditions used in RNA analyses were harvested and grinded with liquid nitrogen and homogenized in 3.5 ml of lysis buffer consisting of 200 mM Tris-HCl (pH 8.5), 50 mM KCl, 25 mM MgCl_2_, 2 mM ethylene glycol tetra acetic acid (EGTA), 2% PTE (10-Tridecyl Polyoxyethylene ether), 1% octylphenoxy poly(ethyleneoxy)ethanol (IGEPAL), 1% Tritón X-100, 0.1% DTT (1M), 0.2 mg/ml cycloheximide, and 0.5 mg/ml Heparin. After the lysis, samples were centrifuged twice at 12,000 g at 4°C for 15 min. After clarification by centrifugation, supernatants were placed on 60% sucrose in gradient buffer containing 50 mM Tris-HCl (pH8.5), 20 mM KCl, 10 mM MgCl_2_, and 0.2 mg/ml cycloheximide and centrifuged in a Beckman 80Ti rotor at 50,000 rpm at 4°C for 3 h. The pellet was resuspended in 400 µl of lysis buffer, loaded onto a continuous 20–60% sucrose gradient, and centrifuged at 45,000 rpm for 2 h in a Beckman SW55Ti rotor. Gradients were fractionated using an ISCO fraction collector with continuous monitoring of absorbance at 260 nm. RNA was isolated from each fraction following the protocol described by (Martinez-Silva *et al*., 2012).

### RNA sequencing and transcriptome analysis

Total RNA was extracted from 7-day-old seedlings of Col-0 and *4ehp-1* grown under control conditions (22°C), after exposure to acclimation at 38°C for 90 min, or after recovery at 22°C and heat stress at 45°C for 2 h each. Three biological replicates were performed for each genotype and condition. RNA quality was assessed using a Bioanalyzer (Agilent), and samples with RIN > 8 were used for library preparation. Strand-specific RNA-seq libraries were prepared using the TruSeq Stranded mRNA Library Prep Kit (Illumina) and sequenced on Illumina HiSeq 2000 analyzer using paired-end reads. Sequencing and library construction were performed by Unidad Universitaria de Secuenciación Masiva de DNA (UUSMD, UNAM-IBT, Mexico).

Raw reads were quality-filtered and trimmed using Trim Galore (v0.6.6). Cleaned reads were mapped to the *Arabidopsis thaliana* TAIR10 reference genome using HISAT2 (v2.2.1). Read counts were generated using HTSeq (v0.13.5) with default parameters. Differential gene expression analysis was performed using the DESeq2 package in Galaxy (Galaxy, 2024). Genes with adjusted *p*-value<0.05 and |log₂FC| >1 were considered differentially expressed. Venn diagrams were constructed with BioVenn (Hulsen *et al*., 2008). Gene Ontology (GO) enrichment analysis was performed using the ShinyGO v0.85 platform (Ge *et al*., 2020) with a false discovery rate (FDR) threshold of 0.05.

### Protein extraction and electrophoretic separation

The same tissues and temperature conditions used in RNA analyses were collected for protein isolation. Total proteins were extracted from 1 g of plant material following the protocol described in (Herrera-Diaz *et al*., 2018). Proteins were dissolved in 0.5 mL of IEF buffer (8M urea, 2M thiourea, 4% (w/v) CHAPS, 2% (v/v) Triton X-100, EDTA-free Complete^TM^ protease inhibitors (Roche Molecular Diagnostics) and 50 mM dithiothreitol). Samples were cleared by 20 min centrifugation at 18,000 *g*. Concentration was measured in a Nanodrop and corrected with respect to a standard by densitometry of denaturing gel electrophoresis (SDS-PAGE). For 2D gel separation, 300 µg of purified protein sample was adjusted to 250 μL with IEF and used to hydrate 11 cm IPG strip (p*I* 3-10) in a Protean IEF cell unit (BioRad). Following isoelectrofocusing, strips were incubated in equilibration buffer (1.5 M Tris-HCl, 6M urea, 30% [v/v] glycerol, 5% [w/v] SDS and 2% (w/v) dithiothreitol) for 15 min and treated with 2.5% (w/v) iodoacetamide dissolved in the same buffer. Second dimension separation was done on 12% SDS-PAGE. Gels were fixed with 50% methanol for 30 min and stained with Coomassie Colloidal (20% [v/v] ethanol, 1.6% [v/v] phosphoric acid, 8% [w/v] ammonium sulfate, 0.08% [w/v] Coomassie Brilliant Blue G- 250) for at least 16 h. Gel images were acquired with the ChemiDoc MP system and analyzed with PDQuest 2D software version 7.0 (BioRad). Spots that met the criterion of reproducible changes between Col-0 and *4ehp-1* at each temperature or between two temperatures were selected for protein identification from paired gels.

### Mass spectrometry protein identification

For shotgun proteomic analyses 5 μg of cleared protein sample was reduced with 50 mM DTT for 20 min and alkylated with 30 mM iodoacetamide for 30 min in the dark. Alkylated samples were purified with Zip Tip C4 pipet tip (Millipore), eluted with 0.1% formic acid (FA) in 4% acetonitrile (ACN) and dried under vacuum. Digestion was performed with 1μg/μL Trypsin Gold (Promega) in 0.05 M ammonium bicarbonate for 16 h at 37°C. The resulting peptides were completely dried under vacuum, rehydrated in 0.1 % FA in 4% ACN and purified by ZipTipU-C18 (Millipore). Mass spectrometry analysis was performed with Synapt G2 high-definition mass spectrometer (Waters Corporation) equipped with a NanoLockSpray ion source (nanoAcquity UPLC Trap Column 5 μm 180 μm ×20 mm), coupled online to a nanoAcquity UPLC Column 1.8 μm, HSST3 C18, 75 μm ×150 mm. The global ProteinLynx version 2.4 server (PLGS; Waters Corporation) and *Arabidopsis thaliana* protein database TAIR 10.1 downloaded from NCBI were used for spectrometric data analysis. PLGS scores with confidence > 95 % were accepted as correct.

For 2D-PAGE differential spot identification, excised gel spots were individually cut into small (∼1 x 1 mm^2^) pieces with a scalpel. Pieces were distained, reduced, alkylated, and digested with trypsin as previously described (Herrera-Diaz *et al*., 2018). Peptides were completely dried under vacuum, rehydrated in 0.1% FA, purified by ZipTipU-C18 (Millipore), dried again, and stored at −80°C until analysis. Peptides were reconstituted in 15 μL of 0.1% FA, 5% ACN and the whole volume was loaded into a Thermo UltiMate 3000 HPLC system with a column Acclaim PepMap RSLC C18 (75 μm × 15 cm). A constant flow of 250 nL/min of a mix of 0.1% (v/v) FA in water (Buffer A), and 0.1% (v/v) FA in HPLC grade ACN (Buffer B) was used in a linear gradient of 50 min from 1 to 50% B. Electrospray ionization (ESI) and m/z detection were made with a Q-TOF mass spectrometer (Impact II, Bruker). Calibration was performed every six injections using the ESI-TOF Tuning mix (Agilent). Protein identifications were made processing the raw files with the Compass Data Analysis software (version 4.2 SR2, Bruker) and Mascot 2.4.1 (Matrix Science): trypsin as the digestion enzyme, one miscleavage allowed, carbamidomethyl Cys as a fixed modification and oxidation on Met as variable modification. Monoisotopic peptide masses were searched with 1.2 Da peptide mass tolerance and 0.6 Da fragment mass tolerance. FDR was set to 1% with the percolator and peptide decoy options active. The reviewed and unreviewed UniProt databases for *Arabidopsis thaliana* were used. Proteins with Mascot score > 16 and detection of at least two significant peptides, were considered as successful identifications.

## Results

### The *4ehp-1* mutant displays mild developmental phenotype affecting the flowering timing

To investigate the role of 4EHP in *Arabidopsis thaliana*, we used a T-DNA insertional mutant line SALK_131503 (Fig. 1a) derived from the ABRC T-DNA insertion collection (Alonso *et al*., 2003), referred here as *4ehp-1*. This line has been previously employed in other reports (Keima *et al*., 2017; Lin *et al*., 2025). Its homozygosity was confirmed by PCR genotyping with the appropriate primers (**Fig. 1b, Supporting Information Table S1**). Compared to wild-type Col-0 plants, the mutant did not show significant transcript level changes in *eIF4E1*, *eIF(iso)4E*, or genes adjacent to *4EHP* in chromosome 5, *APRL7* and *CSD3*, while full length *4EHP* transcript was absent (**Supporting Information Fig. S1a**). In addition, we generated three independent stable transgenic lines expressing *4EHP-GFP* under the control of the CaMV 35S promoter in the *4ehp-1* background (*4ehp-1/C.B*, *4ehp-1/C.C*, and *4ehp-1/C.D*). RT-qPCR analysis confirmed overexpression of the *4EHP* transgene in all three complemented lines (**Fig. 1c**).

**Figure 1.**
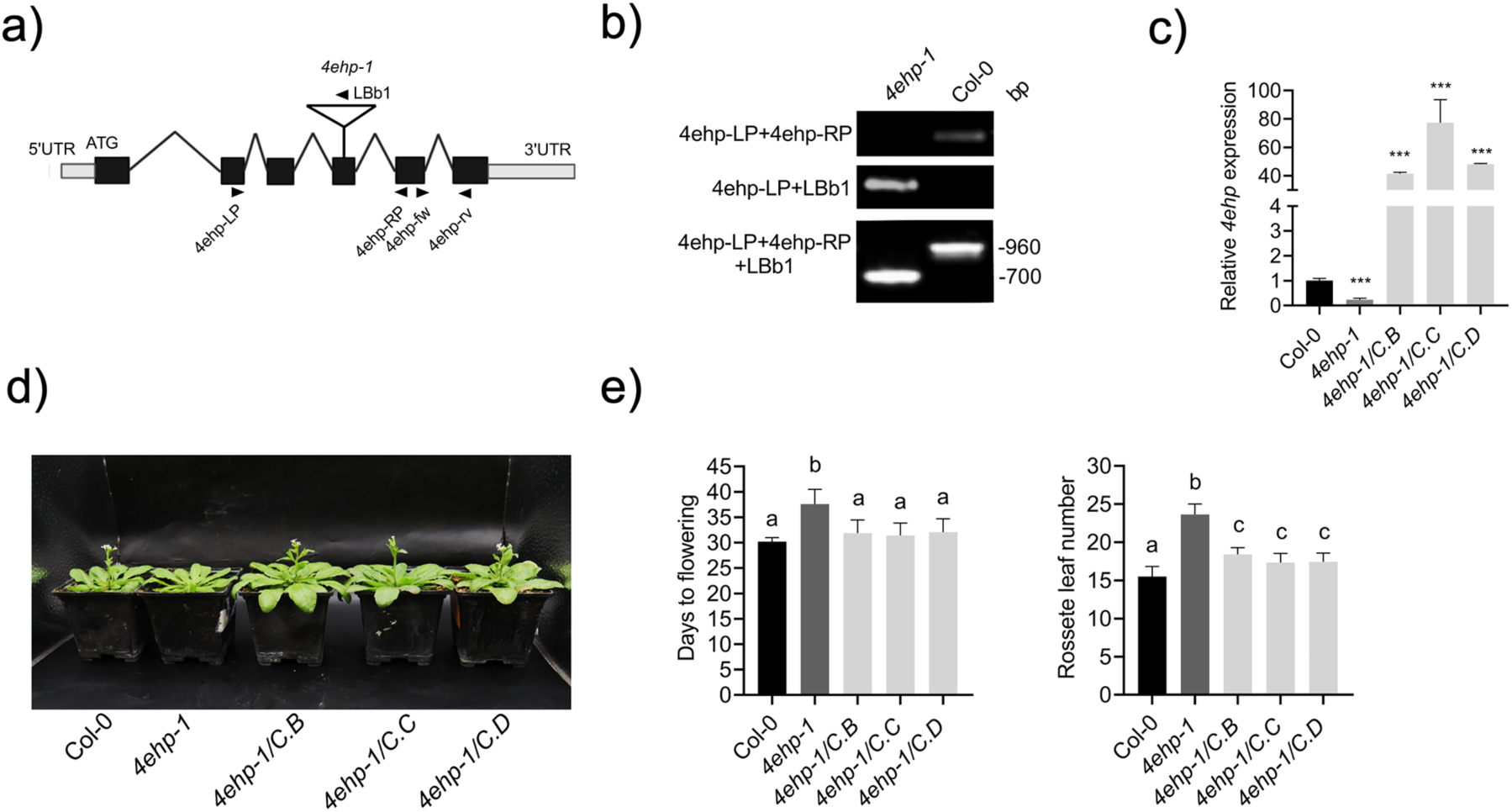
Characterization of the *4ehp-1* mutant line. **a)** Schematic representation of the *4EHP* gene (At5g18110) structure and T-DNA insertion (SALK_131503). Exons are shown as black boxes, introns as connecting lines, 5’ and 3’ untranslated regions (UTRs) as gray boxes. Position of the T-DNA insertion is indicated by an inverted white triangle. Primers used for genotyping (RP, LP border T-DNA insertion, and LBb1 within the T-DNA) and RT-qPCR (4ehp-fw, 4ehp-rv) are indicated by black arrows. **b)** End-point PCR genotyping shows that *4ehp-1* plants are homozygous for the T-DNA insertion. A combination of three primers was used to detect both the fragment corresponding to wild type Col-0 and T-DNA insertion in *4ehp-1*, according to the map shown in a). **c)** *4EHP* mRNA accumulation in Col-0, *4ehp-1*, and three independent mutant lines complemented with p35S:4EHP-GFP (C.B, C.C, and C.D) was analyzed by RT-qPCR. Total RNA was extracted from 7-days-old seedlings grown under long-day conditions at 22°C. RNA levels were normalized to *ACTIN 2* (At3g18780). Data are mean ± SD from three independent biological replicates. Significant difference with respect to Col-0 was calculated by two-tailed Student’s t-test; *** P < 0.001. **d)** Flowering delay for *4ehp-1* under long-day photoperiod. Representative photographs at bolting for Col-0, mutant and the three complemented lines. **e)** Quantitative representation of flowering time as days to bolting (left) or total rosette leaf number at bolting (right) for each genotype. Data represent mean ± SD from at least 20 individual plants per genotype across three independent experiments. Different letters indicate statistically significant differences (one-way ANOVA with Tukey’s post-hoc test, P < 0.05).

The most evident developmental phenotype observed in *4ehp-1* plants was a significant delay in flowering time (**Fig. 1d**). Under LD conditions, *4ehp-1* plants bolted on average one week later than WT (Col-0) and produced more rosette leaves at bolting (**Fig. 1e**). Otherwise, soil germination, seedling growth, leaf, and flower morphologies in the mutant were similar to WT (**Supporting Information Fig. S1b**). Leaf number increase corresponded exclusively to the continued leaf initiation in *4ehp-1* plants due to a delayed reproductive transition. The flowering time and leaf number were rescued in all three complemented lines, confirming that late-flowering phenotype was due to the loss of 4EHP function. Delayed bolting was observed in both LD and SD conditions, indicating that the effect is largely photoperiod independent (**Supporting Information Fig. S1c**).

To test whether 4EHP absence affected cell growth and division, primary root elongation, lateral root primordium formation, and cell proliferation at the root apical meristem were measured (**Supporting Information Fig. S1d**). None of these tissues displayed significant differences compared to WT, at least for the first ten days after germination. This suggests that 4EHP might specifically influence developmental processes but does not affect general cell division activity.

### Loss of 4EHP confers enhanced thermotolerance

eIF4E family members are involved in translation regulation in response to temperature stress conditions (Desroches Altamirano *et al*., 2024; Wu *et al*., 2024). In addition, Arabidopsis 4EHP was identified in proteomic analysis of stress granules during heat stress (Gutierrez-Beltran *et al*., 2021) and orthologs in other species have been shown to participate in stress responses (Christie & Igreja, 2023). Hence, we examined whether *4ehp-1* displayed altered responses to heat stress. An acquired thermotolerance assay, consisting of priming treatment at 38°C, recovery for 2h at 22°C, and further exposure to heat stress (HS) at 45°C, was applied to seven days-old seedlings. The *hsp101* null mutant (*hot-1*) was included as highly susceptible control. Phenotypes were registered nine days after the treatment, by counting the number of seedlings with severe, moderate, and low damage, according to the number of expanded green leaves. More than 50% of Col-0 seedlings showed severe damage. In contrast, *4ehp-1* seedlings recovered significantly better, with 40% of plants presenting low damage and only around 25% with severe damage (**Fig. 2a**, right panel). The Col-0 phenotype was restored in three complemented lines, indicating that enhanced thermotolerance was attributable to 4EHP loss. In contrast to *4ehp-1*, the acquired thermotolerance response of *eif4e* and *eifiso4e* mutants was comparable to Col-0 (**Supporting Information Fig. S2a**).

**Figure 2.**
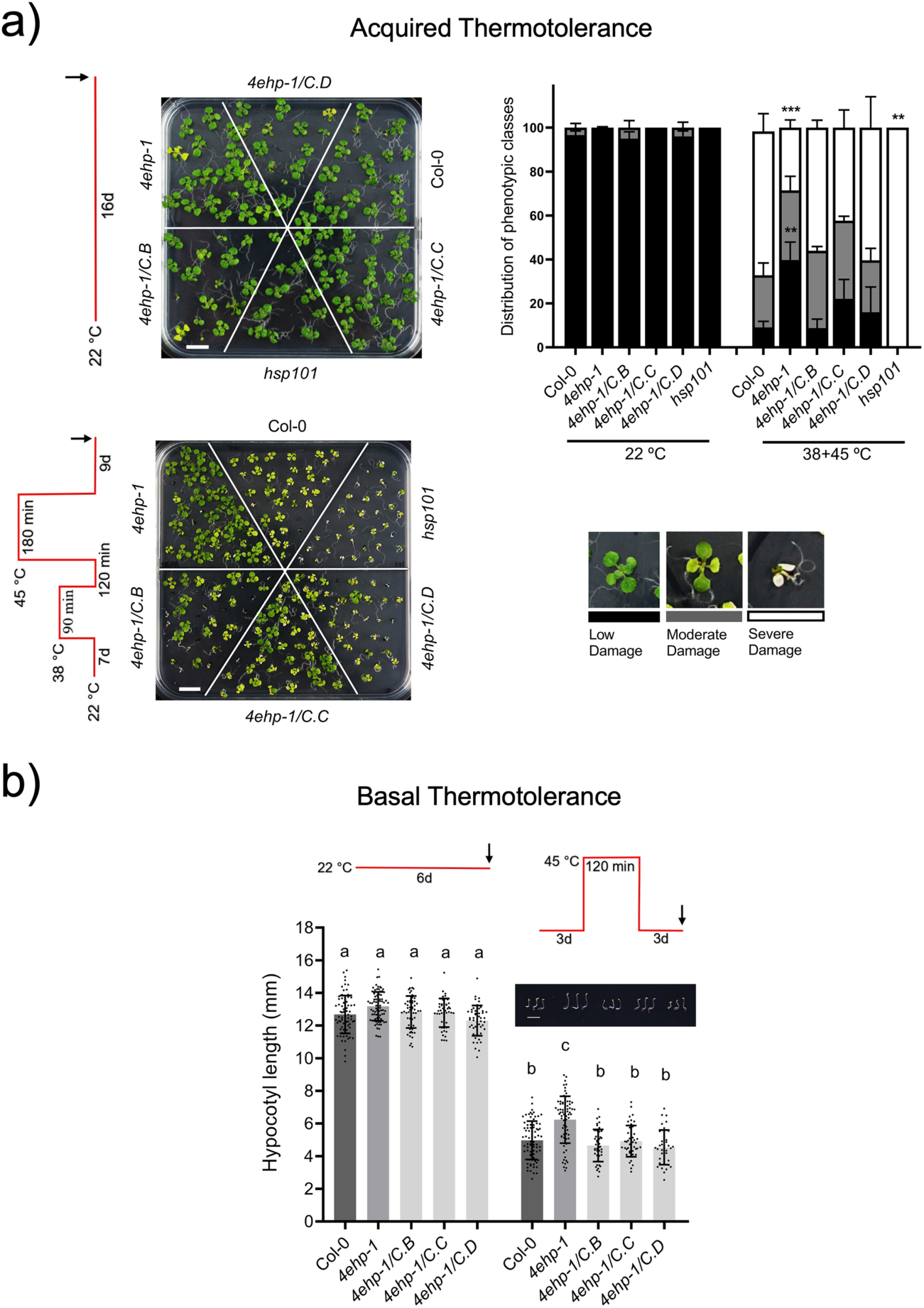
The *4ehp-1* mutant exhibits enhanced thermotolerance. **a)** Acquired thermotolerance evaluated by the level of damage upon recovery from heat stress (HS) applied to acclimated seedlings. Seven-days-old seedlings were incubated at 38°C for 90 min, followed by a recovery period of 120 min at 22°C and heat shock (HS) at 45°C for 180 min. The seedling phenotype was registered nine days after the heat treatment (right panel) and distributed in one of three phenotypic classes: low (black bars), moderate (grey bars), or severe (white bars) damage by counting the number of expanded green leaves, including cotyledons. Control seedlings remained at 22 °C for the same time (16 days). Data represent mean ± SD from 100–160 seedlings per genotype divided into five biological replicates. Asterisks indicate significant differences compared with Col-0 (two-tailed Student’s t-test; *** P < 0.001, ** P < 0.01). **b)** Basal thermotolerance evaluated by hypocotyl elongation. Three-day-old seedlings grown under dark conditions were incubated at 45°C for 120 min. Three days after heat treatment, hypocotyl length was measured in control and HS. Control seedlings remained under dark at 22 °C for 6 days. Data represent mean ± SEM from 50–100 hypocotyls per genotype divided into five biological replicates. Different letters indicate statistically significant differences (one-way ANOVA with Tukey’s post-hoc test, P < 0.05). Representative photographs of seedlings 3 days after HS are shown inside the graph.

To assess whether *4ehp-1* plants also possess superior basal thermotolerance (intrinsic heat tolerance without prior acclimation), we exposed three days-old, dark-grown seedlings to 45°C for two hours and measured hypocotyl elongation after three days of recovery in the dark. In this experiment, *4ehp-1* seedlings showed significantly greater hypocotyl growth following stress compared to Col-0 or the complemented lines, which was not attributable to any difference between lines at the time of HS application or their growth under control conditions (**Fig. 2b**; **Supporting Information Fig. S2b**). Altogether, the thermotolerance assays revealed that the *4ehp-1* mutant exhibited a reduced sensitivity to HS, indicating that this class II eIF4E protein might interfere with growth during thermal stress adaptation.

### 4EHP re-localizes to cytoplasmic stress granules upon heat stress

To investigate the molecular basis of enhanced thermotolerance in *4ehp-1*, we examined the subcellular localization of 4EHP-GFP during HS. Under control growth conditions, 4EHP-GFP showed diffuse cytoplasmic distribution in both, transiently transformed protoplasts overexpressing 4EHP-GFP, and the stable complemented lines (**Fig. 3a, b**). Upon HS at 40°C for 35 minutes, 4EHP-GFP accumulated into discrete cytoplasmic foci in the protoplasts, with an average of 40 foci per cell (**Fig. 3a**). In five days-old root tips of the complemented lines, 4EHP-GFP also localized to cytoplasmic foci under HS (**Fig. 3b**). Such re-localization was reversible, since it returned to a diffuse pattern following a recovery at 22°C for 6 hours. Similar distribution dynamics upon HS was observed for a construct under the *4EHP* promoter (p4EHP:4EHP-GFP) in either transiently transformed protoplasts or root tips of stable mutant lines (**Supporting Information, Fig. S3**). The presence of 4EHP in cytoplasmic condensates was also detected after incubation at 38°C for 1.5 h (**Supporting Information, Fig. S3**).

**Figure 3.**
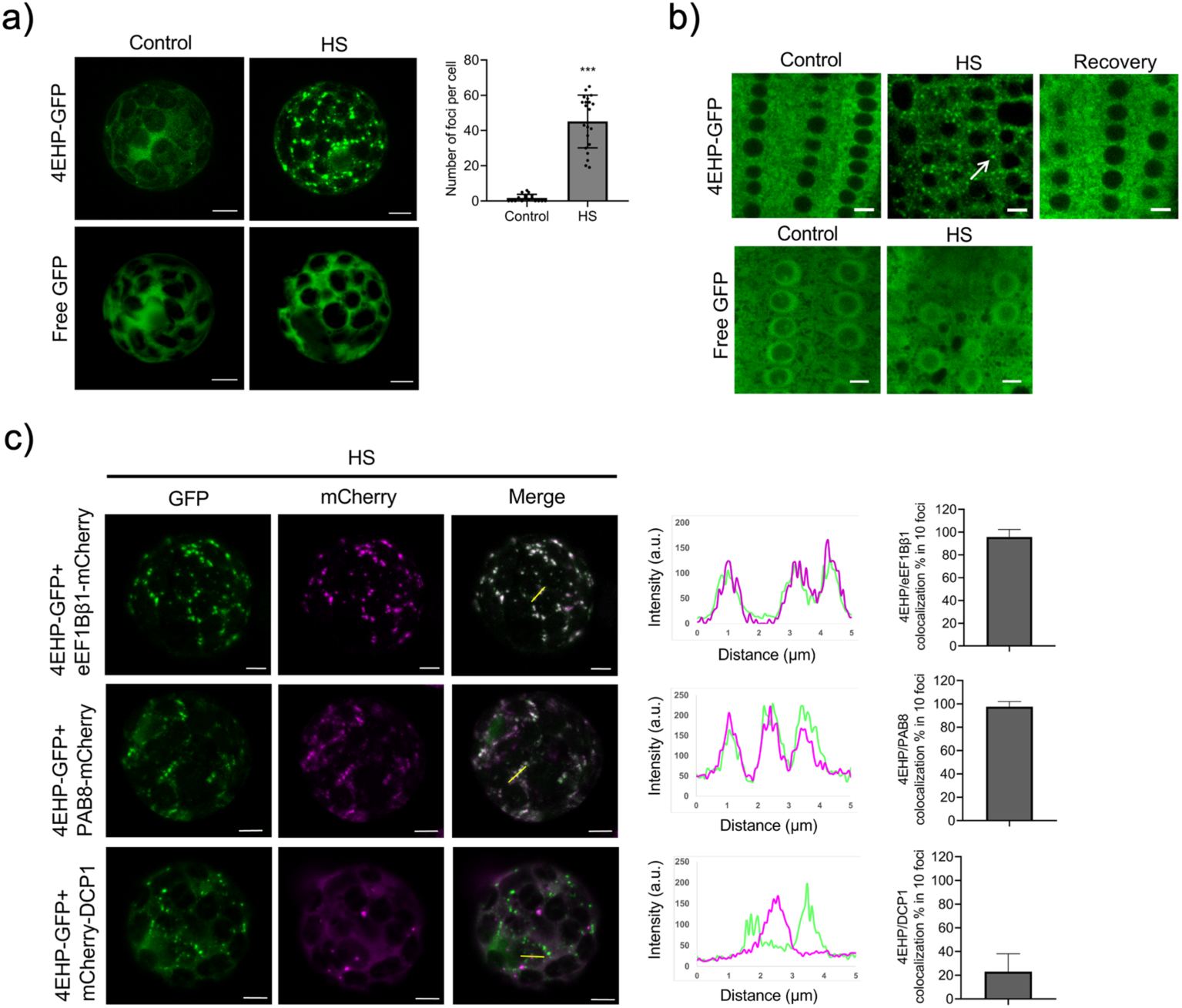
The 4EHP protein is recruited to stress granules during HS. **a)** Protoplasts transiently expressing p35S:4EHP-GFP present different fluorescence distribution after HS at 40°C for 35 min compared to control conditions or free GFP. Quantification of the number of fluorescent foci per cell in control and HS protoplasts is shown at right. Asterisks indicate significant difference between treatments n=22 (two-tailed Student’s t-test; *** P < 0.001). Scale bar = 5 μm. **b)** Localization of 4EHP-GFP in root cells of five-day-old p35S:4EHP-GFP complemented mutant line. Seedlings were incubated at 22°C (control) or at 40°C for 35 min (HS) with a recovery period of 6 h at 22 °C. White arrow indicates 4EHP-GFP accumulation in cytoplasmic foci. Scale bar = 5 μm. **c)** Confocal images of protoplasts transiently co-expressing 4EHP-GFP with stress granule (SG) markers: eEF1Bβ-mCherry, PAB8-mCherry, or processing body (PB) marker: mCherry-DCP1 after HS at 40°C for 35 min. All reporters were under the 35S CaMV promoter. Fluorescence intensity plots of GFP (green) and mCherry (magenta) are shown for the yellow line indicated on merged images (center). Co-localization between 4EHP and the indicated marker is indicated as percentage per 10 foci for n=15-30 independent protoplasts (right graphs). Scale bar = 5 μm.

Reports on 4EHP animal orthologues have indicated that the protein localizes to stress granules (SGs) or processing bodies (PBs) under stress conditions to regulate the fate of specific mRNAs (Christie & Igreja, 2023). To explore the identity of HS-induced 4EHP-GFP foci detected in Arabidopsis cells, we performed co-localization experiments with SG and PB markers (**Fig. 3c**). Co-expression of 4EHP-GFP with mCherry-tagged eEF1Bβ and PAB8 (SG) showed extensive overlap in the HS induced foci, approaching 100% of co-localization. In contrast, co-localization with the PB marker DCP1 was significantly lower, with only approximately 20% of 4EHP-GFP foci overlapping with the DCP1 signal (**Fig. 3c**, right panel). Fluorescence intensity profiles across individual granules confirmed coincident peaks between 4EHP and SG markers, whereas 4EHP and DCP1 signals were mainly separated (**Fig. 3c**, middle panel). These findings support that Arabidopsis 4EHP is preferentially recruited to SGs under HS conditions.

### The absence of 4EHP affects transcriptome profiles during heat stress (HS)

As cap-binding protein, 4EHP is expected to influence the fate or stability of mRNAs (Christie & Igreja, 2023). Therefore, we evaluated whether its absence affected overall transcript levels under normal and high temperature conditions, where samples were taken just after incubation at the indicated temperatures (**Fig. 4a**) – control (22 °C), acclimated for 1.5 h (38 °C), recovered and heat-stressed for 2 h (38+45 °C). Transcriptomic analyses were done for three independent biological replicates. mRNA levels were highly correlated across the biological replicates and differed between temperatures and lines (**Supporting Information Fig. S4**). Transcripts belonging to differentially expressed genes (DEGs) were considered if they displayed significant (adjusted p-value<0.05), and at least two-fold change (FC) between compared samples (absolute log2FC>1) in our comparisons.

**Figure 4.**
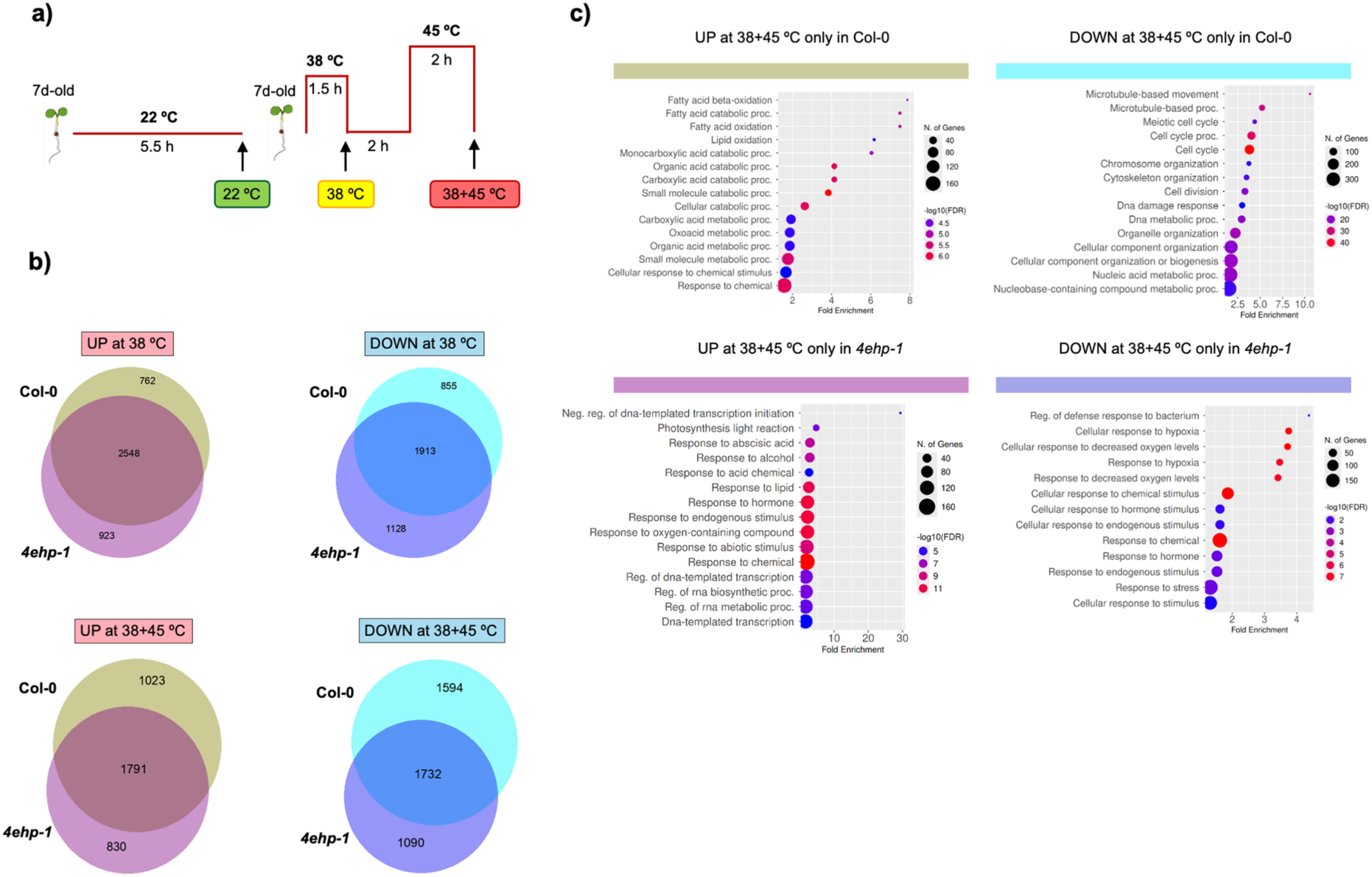
Transcriptomic responses to heat stress in *4ehp-1* and Col-0. a) Schematic overview of the acquired thermotolerance assay used in transcriptome analyses. b) Venn diagrams showing overlapping or differential transcriptional changes during acclimation and HS for Col-0 and *4ehp-1*. Fold change (FC) was calculated as reads detected at 38 °C or 38+45 °C versus reads detected at 22 °C for the same line. UP, up-regulated genes (log2FC>1, FDR<0.05); DOWN, down-regulated genes (log2FC<1, FDR < 0.05). c) Biological Process (BP) enrichment is shown for unique significant changes for Col-0 and *4ehp-1* below each Venn diagram. The rectangle colors match the Venn diagram portion corresponding to each class of unique changes.

First, we analyzed the effect of temperature changes separately on Col-0 and *4ehp-1* transcriptome profiles and compared the observed DEGs with respect to control (Fig. 4). During the acclimation stage (38 °C) we found great overlap between the DEGs in wild type and *4ehp* mutant line; 2548 coincident transcripts were upregulated and 1913 downregulated (**Fig. 4b, Supporting information Table S2**). In addition, 762 transcripts appeared upregulated in Col-0 and 923 in the mutant only, while 855 decreased in Col-0, and 1128 in *4ehp-1*. After recovery and HS (38+45 °C), the number of distinctive DEGs was greater for Col-0 (**Fig. 4b**). Genes significantly up-regulated at 45 °C only in Col-0 (1023) were related to fatty acid and organic acid catabolic biological processes, whereas those downregulated (1594) were significantly enriched in cell cycle and division, DNA metabolism and chromosome organization (**Fig. 4c**). On the other hand, *4ehp-1* specific DEGs upon HS, albeit less in number (830 upregulated and 1090 downregulated), were represented by responses to stress stimulus, hormone-related, oxygen levels, and RNA metabolism.

Next, we searched for transcriptome changes due to the absence of 4EHP under each temperature condition: 22 °C, 38 °C and 38+45 °C (**Fig. 5; Supporting Information Table S3**). Under control conditions, we found 495 DEGs in *4ehp-1* mutant distributed in 257 upregulated and 238 downregulated transcripts (**Fig. 5a**). The number of DEGs increased to 695 at 38 °C and 1877 after heat stress (38+45 °C), with most DEGs in the last condition corresponding to upregulated transcripts (1240). Importantly, we observed between the 20 most significant upregulated DEGs in *4ehp-1* at 38+45 °C several transcripts coding for heat stress proteins, such as *HSP90.1* (At5g52640), *HSP22* (At4g10250), *HSP21* (At4g27670), *HSP17.6II* (At5g12020). The *HSP22*, *HSP 17.6II* and other HSP coding transcripts were also identified between the significant DEGs at control temperature.

**Figure 5.**
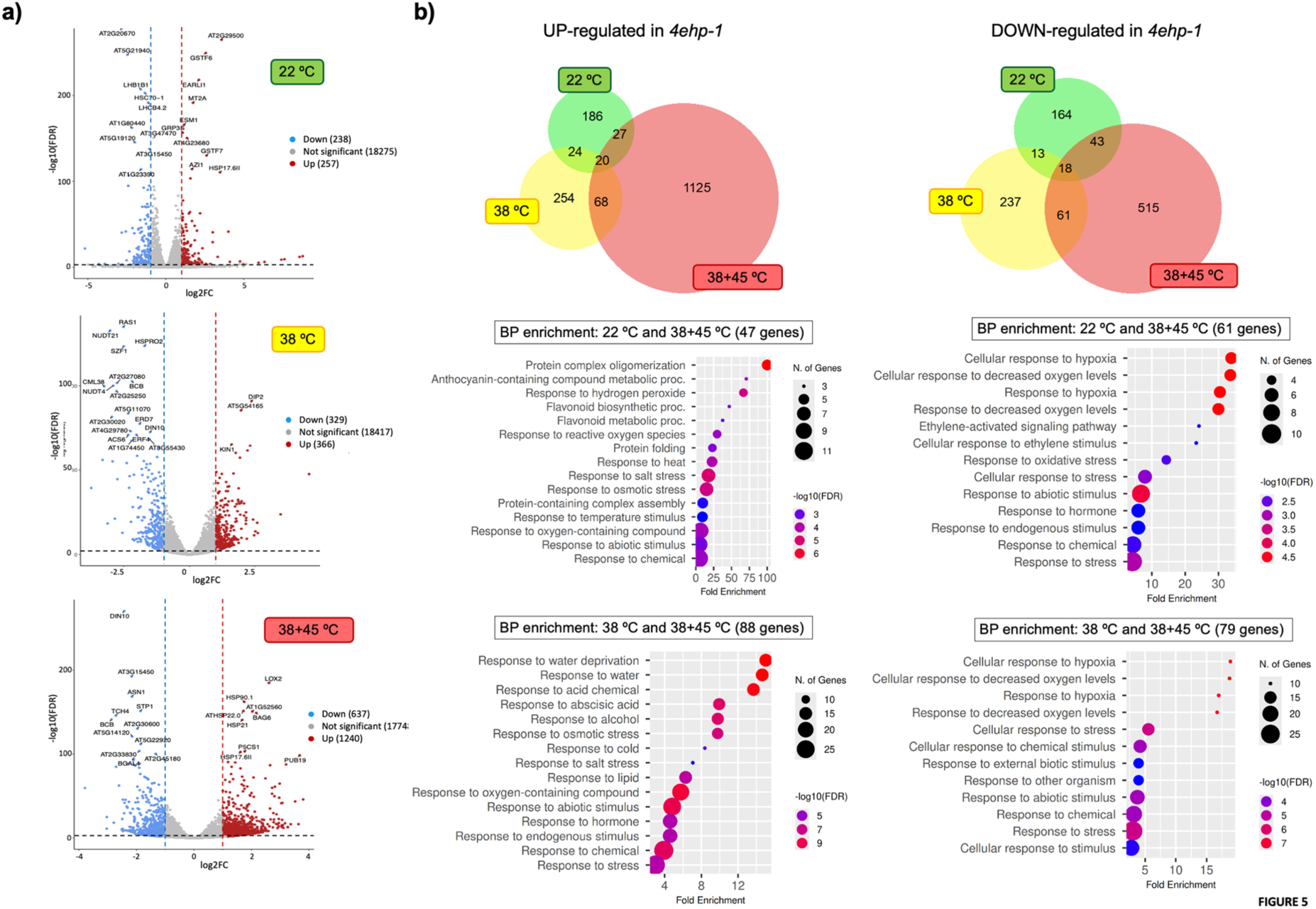
Transcriptome comparisons between *4ehp-1* and Col-0. a) Transcriptional changes in the absence of 4EHP under control (22 °C), acclimation (38 °C) and heat stress (38+45 °C) temperatures. Fold change (FC) was calculated as reads detected in *4ehp-1* versus reads detected in Col-0 at each temperature. Volcano plots show significant DEGs in color (absolute log2FC > 1 and FDR < 0.05). Down-regulated genes (blue dots); up-regulated genes (red dots). b) Venn diagrams of upregulated and downregulated genes in 4ehp-1 mutant among the three temperatures. The GO enrichment results for “Biological Process” of co-upregulated and co-downregulated *4ehp-1* genes between 22 and 38+45 °C or between 38 and 38+45 °C are shown below the Venn diagrams. The number of genes is indicated in parenthesis. Plots were generated in ShinyGO 0.85, with FDR < 0.05.

Venn diagram analysis for mutant to WT comparisons showed 115 upregulated and 122 downregulated DEGs, overlapping among temperatures (**Fig. 5b**). Between them, 47 upregulated DEGs were shared at 22 and 38+45 °C. This group was enriched in stress regulators, including protein oligomerization and HS, characteristic to HSPs. Similarly, the 88 upregulated DEGs corresponding to 38 °C and 38+45 °C overlap included significant enrichment of stress-related GOs; albeit water-deprivation stress appeared as highlighted in this group. Interestingly, the *4ehp-1* downregulated DEG overlaps for the same temperatures were enriched in hypoxia response, ethylene signaling or biotic stress GOs, different from categories observed in the upregulated transcripts.

The highest number of upregulated DEGs in *4ehp-1* compared to Col-0 was observed upon HS. Enrichment analysis for these DEGs indicated microtubule movement, chromatin organization, response to water deprivation and cell cycle as the most significant biological processes for this group (**Supporting Information Fig. S5**). The cellular component and molecular function GO enrichment was consistent with microtubule-based movement, nuclear activity, and DNA metabolism significantly higher in the mutant at 38+45 °C, with respect to WT. On the other hand, downregulated DEGs were enriched in hypoxia and decreased oxygen level responses, with genes coding for cell wall constituents, transmembrane transporters, and peroxidase activity.

Notably, overall transcriptome profiling suggested that in the absence of 4EHP, heat stress response and cell cycle activity are enhanced with respect to Col-0 after HS. This is in line with our phenotype analysis and indicates that the mutant line is insensitive to increased temperature stress and thus displays relatively unaffected growth compared to Col-0 (**Fig. 2a**).

### *HSP* transcripts over-accumulate and remain translated under HS in the absence of 4EHP

Transcriptome profiles together with cytoplasmic co-localization of Arabidopsis 4EHP in SGs upon HS are suggestive of a role for this protein in mRNA stability or translation. Since HSPs were found between the most significant transcripts changing their accumulation in the *4ehp-1* mutant, we decided to evaluate the RNA level and polyribosomal distribution of four relevant HSPs by RT-qPCR under each temperature condition. HSP101 (At1g74310), HSP70B (At1g16030), Class II HSP17.6A (At5g12030), also termed HSP17.7, and Class I HSP17.6C (At1g53540) act as chaperones at different stages of protein homeostasis under heat stress (see Supporting Information Table S4 for different HSPs in *Arabidopsis thaliana*). Their total RNA level was highly induced during acclimation (38°C) in both Col-0 and the *4ehp-1* mutant (**Fig. 6a**). However, at control temperature (22°C) *HSP101*, *HSP17.6A* and *HSP17.6C* displayed significantly higher expression levels in the mutant. Significant upregulation of all four mRNAs was also observed in *4ehp-1* compared to Col-0 in HS conditions (38+45°C). Such upregulation was abolished in the complemented line *4ehp-1*/C.C (**Supporting Information Fig. S6a**). According to this, the absence of 4EHP is responsible for an overaccumulation of several *HSP* transcripts in control and heat stress temperatures, which might in turn contribute to a higher thermotolerance of the mutant.

**Figure 6.**
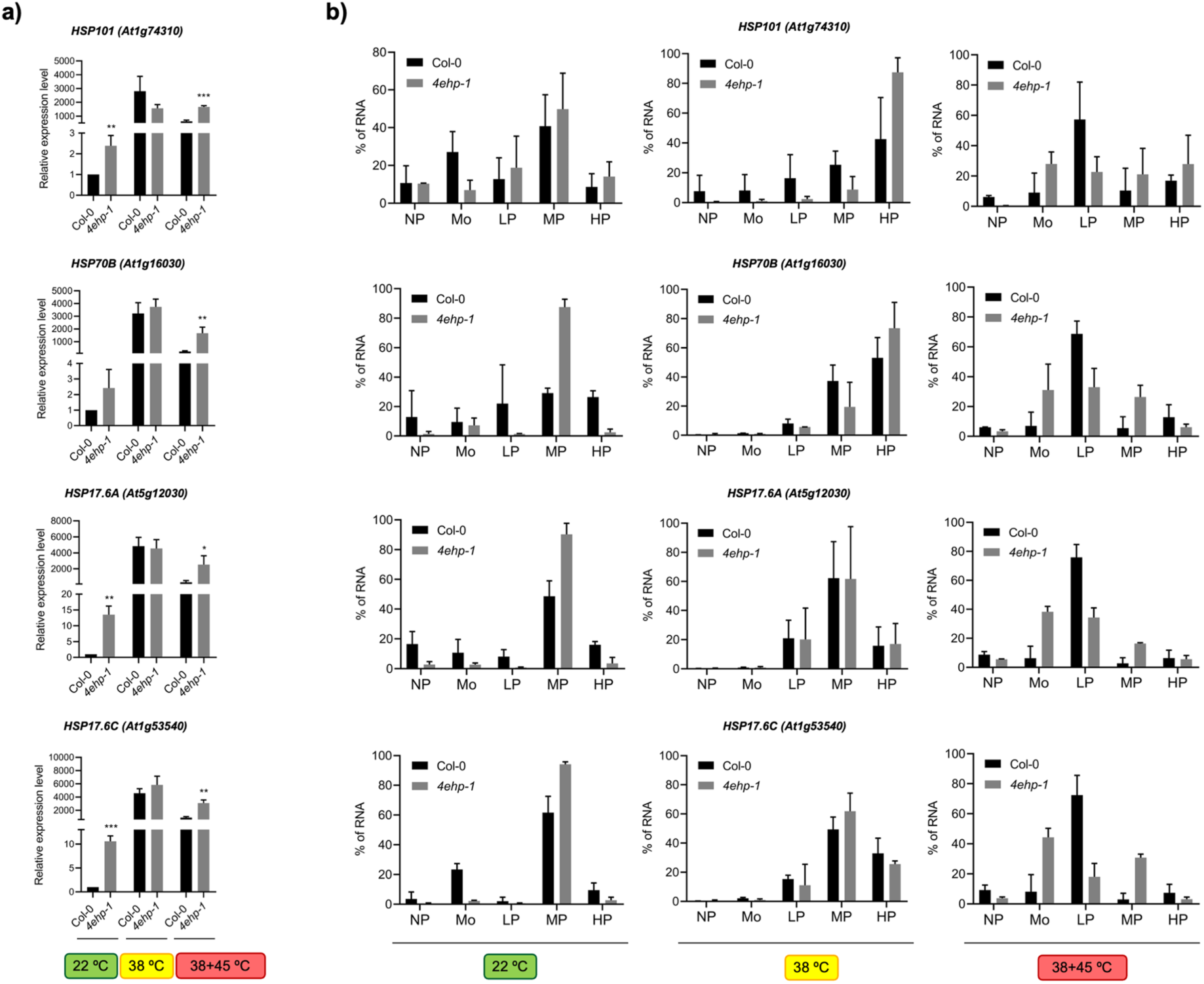
HSP transcripts accumulate at a higher level in the *4ehp-1* and are actively translated under high temperature conditions. a) Four transcripts coding for heat shock proteins, *HSP101*, *HSP70B*, *HSP17.6A* and *HSP17.6C*, were analyzed at total RNA levels in Col-0 and *4ehp-1* under control temperature (22°C), acclimation (38°C) and heat stress (38+45°C) by RT-qPCR. Relative expression levels represent the abundance with respect to Col-0 at 22°C normalized by *ACTIN2*. Data are the mean of three independent biological replicates with three technical replicates each. b) The same transcripts and temperatures shown in (a) were evaluated on pooled fractions of polyribosome profiles. Each profile was composed of 17 fractions, which were pooled according to the absorbance registered at 260 nm and rRNA patterns on agarose gels (Supporting information, Fig. S6): NP (non-polysomes), Mo (monosomes), LP (low polysomes), MP (middle polysomes), HP (high polysomes). RNA was isolated from the pooled fractions and subjected to RT-qPCR. The fraction with highest Ct value was used as reference for 2^DCt^ calculation, and the RNA level was represented as percentage from the sum of all fractions. Error bars correspond to ±SD of two biological replicates with three technical replicates each.

At the level of *HSP* mRNA translation, an overall similar distribution along polyribosomes was found for Col-0 and *4ehp-1* at each temperature. An increase in recruitment towards high polyribosomes was observed for *HSP101* and *HSP70B* at 38°C, suggesting more efficient translation, when compared to 22°C (**Fig. 6b**). This pattern differed from what was seen for a control mRNA *ACTIN2*, which displayed few changes at 38°C (**Supporting Information Fig. S6c**). *HSP17.6A* and *HSP17.6C* distribution also remained unchanged between 22 and 38°C, suggesting similar translation efficiency at both temperatures. Transcript patterns along polyribosomes showed important changes upon heat stress (38+45°C). In Col-0, a major peak was detected in low polyribosomes for all four *HSP* mRNAs translation, while in *4ehp-1* their distribution was similar for monosomes, low polysomes and middle polysomes (**Fig. 6b**; right panels). In regards of HSP101, around 20% of mRNA was still detected in high polyribosomes, indicating active translation despite overall translation inhibition upon heat stress (see **Supporting Information Fig. S6b and S6c**). The differences between polyribosomal patterns of *HSP* mRNAs in *4ehp-1* and Col-0 under HS (38+45°C) suggest that 4EHP might contribute to some extent to translation regulation under heat stress. However, the accumulation found at total RNA levels in the mutant could also account for a more homogeneous distribution between different polyribosomal fractions.

### The absence of 4EHP is associated with greater HSP diversity after HS on a protein level

We next evaluated whether the *4ehp-1* mutant presented differences in accumulated proteins at the same temperature conditions used for total and polyribosomal RNA studies. Samples were analyzed by label-free shotgun proteomics, two-dimensional gel electrophoresis (2D-PAGE) and differential spot identification. The overall results are shown in **Fig. 7** and **Supporting information Fig. S7**, **Tables S5** and **S6**. Interestingly, while a greater number of HSPs were identified by shotgun proteomic analysis with 95% confidence for Col-0 rather than for *4ehp-1* at 22 and 38 °C, the opposite was observed after HS (38+45°C). According to the total number of IDs in each sample (**Supporting Information Table S5**), the percentage of identified HSPs was also greater for the mutant after HS, as indicated in the graph of Fig. 7a. Moreover, several of the HSPs identified after HS in *4ehp-1* but not in Col-0 proteomics corresponded to differentially accumulated transcripts (**Supporting information, Table S4**). These included HSP101 (At1g74310), HSP70B (At1g16030), Class II HSP17.6B (At5G12020), and HSP70.11 (At5g28540).

**Figure 7.**
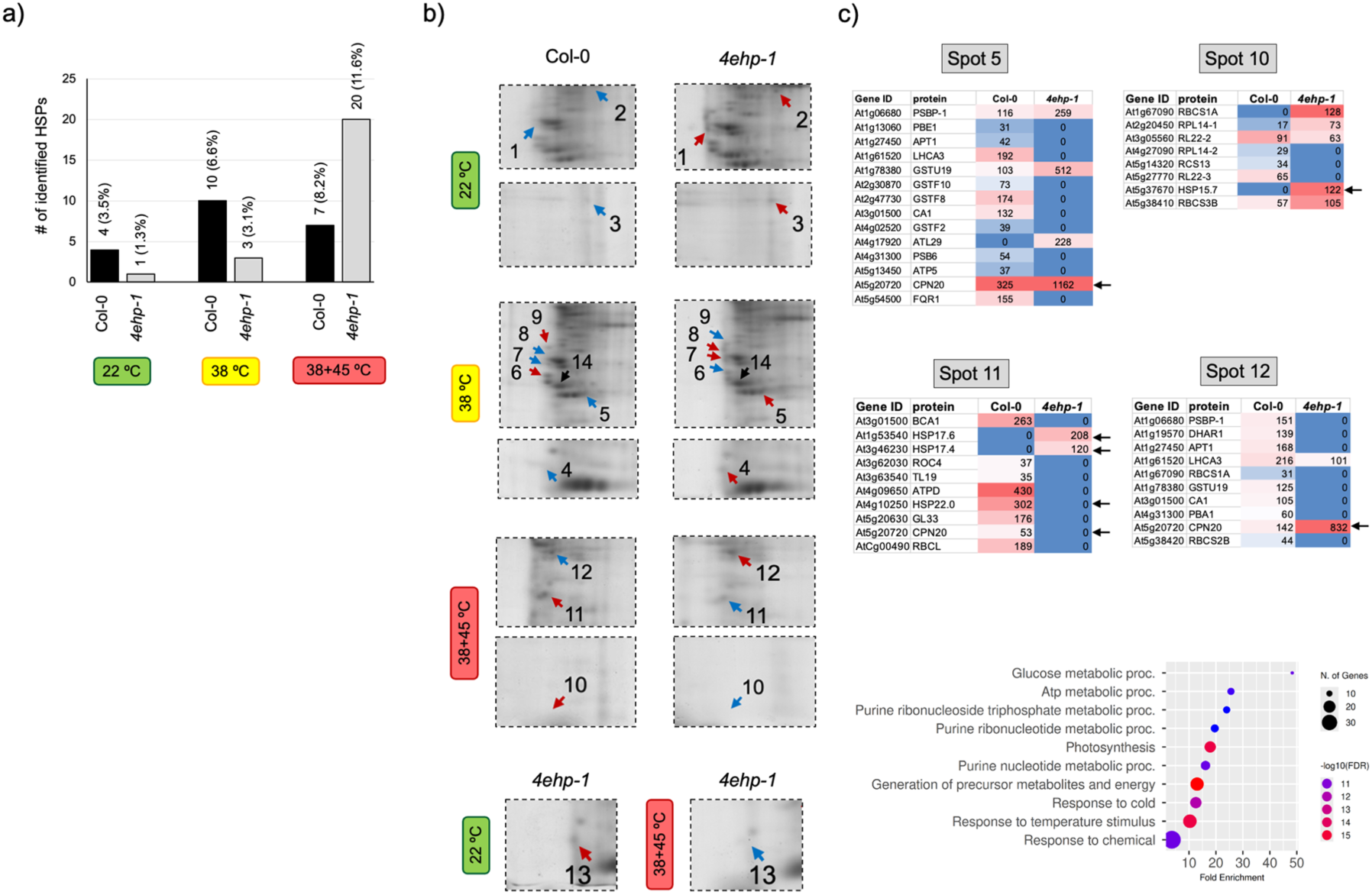
Shotgun and 2D-PAGE protein identifications support differences in HSP distribution between Col-0 and *4ehp-1* upon heat stress. a) Number of HSP proteins identified by shotgun proteomic analysis in Col-0 and the *4ehp-1* mutant at 22°C, 38°C and 38+45°C. The percentage of identified HSPs, according to the total number of proteins identified with 95% confidence, is indicated in parenthesis. The full set of data is available in Supplementary Table S5. b) Differential protein spots between Col-0 and *4ehp-1* at 22°C, 38°C and 38+45°C from 2D-PAGE separation. Each spot is highlighted by arrow (red, increased; blue, decreased). At the bottom, a spot comparison between 22°C and 38+45°C temperatures is shown for the mutant line. Full-sized gels and overall spot distribution are available in Supporting information, Fig. S7. c) Proteins identified with at least two significant peptides from selected paired spots (Col-0 and *4ehp-1*) are represented in heat maps according to their identification score. Arrows indicate HSPs. All proteins identified in differential spots are available in Supporting information, Table S6. The Biological Process enrichment of these proteins is shown in the lower panel.

The same protein samples were subjected to 2D-PAGE separation to access specific spot differences between Col-0 and the mutant (**Supporting Information Fig. S7**). A close-up on gel regions of interest is shown in **Fig. 7b**. Fourteen spots showing differential intensity between lines were mirrored for gels corresponding to each line, excised, and treated for protein identification by LC-MS/MS. The results of all confident identifications (at least 2 significant peptides) are available in **Table S6**. Several proteins were identified by spot for each line, and occasionally the same protein was identified in different spots. This is not unusual, since proteins can migrate at different points in 2D-PAGE depending on post-translational modifications. HSPs were identified in spots 5 (38 °C), 10, 11 and 12 (38+45 °C), all corresponding to comparisons between lines (**Fig. 7c**). According to their score of identification (red to blue colors), most HSPs were characteristic of the *4ehp-1* mutant line, except Hsp22.0 (At4g10250). Between them, HSP15.7 (At5g37670), HSP17.4 (At3g46230) and HSP17.6 (HSP17.6C; At1g53540) were previously identified as differentially accumulated transcripts (increased) by the RNA seq data in *4ehp-1* (Supporting Information Table S4). CPN20 (At5g20720) was identified in spots 5 and 12 with higher scores for the mutant, while in spot 11 it was identified only in Col-0. A gene ontology analysis of all proteins identified in differential spots supported the enrichment of temperature stimulus response in the group, in addition to several metabolic processes (**Fig. 7c**, lower panel).

## Discussion

In this study, we described phenotypic characteristics of a 4EHP mutant line and, for the first time, demonstrated a role for this Class II eIF4E family member in heat thermotolerance. This factor is recruited into stress granules upon heat stress in Arabidopsis and regulates the homeostasis of heat-responsive mRNAs. The effect of 4EHP loss on thermotolerance phenotype differs from what was previously reported for *eif4e* and *eifiso4e* mutants (Salazar-Diaz *et al*., 2021; Zhang *et al*., 2024), suggesting a distinct regulatory role within the eIF4E family. On the other hand, the delayed flowering time is also characteristic to the *eif4e* mutant (Zhang *et al*., 2024) and was recently described for *Brassica napus 4ehp* mutants (Wang *et al*., 2025). Our results showed no apparent root defects in *4ehp-1,* which is consistent with 4EHP targets possibly enriched in aerial meristems, where developmental phase transitions and environmental stress responses must be strictly regulated. The flowering phenotype coupled with enhanced thermotolerance also indicates a trade-off between developmental progression and stress response that may reflect evolutionary optimization of resources under changing thermal exposures. One possibility is that loss of 4EHP constitutively activates stress-response programs that antagonize reproductive maturation, as described for crosstalk between flowering time pathways and abiotic stress signaling (Kazan & Lyons, 2016; Preston & Fjellheim, 2022). Hence, it would be relevant to explore how heat stress affects flowering time in the *4ehp-1* line.

The cellular localization of 4EHP-GFP expressed either under constitutive or its own promoter changed between stress and control conditions. Upon HS challenge, it was localized in SGs like other members of the eIF4E family (Weber *et al*., 2008), possibly through its intrinsically disordered N-terminal region (IDR) that is not involved in eIF4E cap-binding interactions (Marcotrigiano *et al*., 1997; Rosettani *et al*., 2007). These motifs can be necessary and/or sufficient to drive protein re-localization into SGs *in vivo* or to promote liquid–liquid phase separation *in vitro* (Lin *et al*., 2015; Protter *et al*., 2018). In addition, 4EHP exhibits a high degree of co-localization with SG markers like eEF1Bβ1 and PAB8 and was previously identified as HS–sensitive interactor of TSN2, a protein that functions as a scaffold for SG assembly in plants (Gutierrez-Beltran *et al*., 2021). The preferential co-localization of 4EHP with SG markers and its lower presence in P-bodies may indicate a function in transient translation inactivation of target transcripts before their transference to P-bodies for degradation. This is in line with previously described models of functional communication between cytoplasmic mRNP aggregates (Decker & Parker, 2012).

The Arabidopsis 4EHP role in RNA regulation has been poorly documented. Initial characterization indicated that it was able to interact with eIF(iso)4G *in vitro*, but poorly supported translation, suggesting a role in specific mRNA sequestration (Ruud *et al*., 1998). More recently, it was shown to interact with EXA1, which was relevant for potyvirus infection in Arabidopsis (Nishikawa *et al*., 2023). Here we found that the 4EHP function in Arabidopsis heat stress appears to be more like that reported for the human class III factor eIF4E3_A rather than for the canonical 4EHP (eIF4E2). While mammalian eIF4E2 localizes to both P-bodies and stress granules and is primarily associated with mRNA decay and translational repression, eIF4E3_A localizes exclusively to stress granules during heat stress (Frydryskova *et al*., 2016). Hence, 4EHP might perform different functions, depending on the cellular state (**Fig. 8**). Under normal growth conditions, it likely functions as a translational repressor of specific mRNAs responsive to stress, as described for its mammalian counterpart (Chapat *et al*., 2017). Its affinity to mRNAs might be dictated by specific sequences and interactors in addition to 5’cap binding (Villaescusa *et al*., 2009; Morita *et al*., 2012). For example, GYF proteins interacting with 4EHP, such as EXA1, could promote the formation of translational repressor complexes as reported for animal counterparts. Upon temperature increase, 4EHP becomes recruited to stress granules, but during acclimation its influence on HSP client mRNAs is overcome by the burst on their transcription and translation, thus allowing for an appropriate stress response. Persistence of HS promotes the 4EHP regulatory role on HSPs, recruiting more transcripts to the SGs to exert translation inhibition and possibly directing some mRNAs to further degradation. After six hours of recovery, 4EHP returns to a diffuse cytoplasmic localization providing a rapid and reversible regulatory switch for *HSP* mRNA translation.

**Figure 8.**
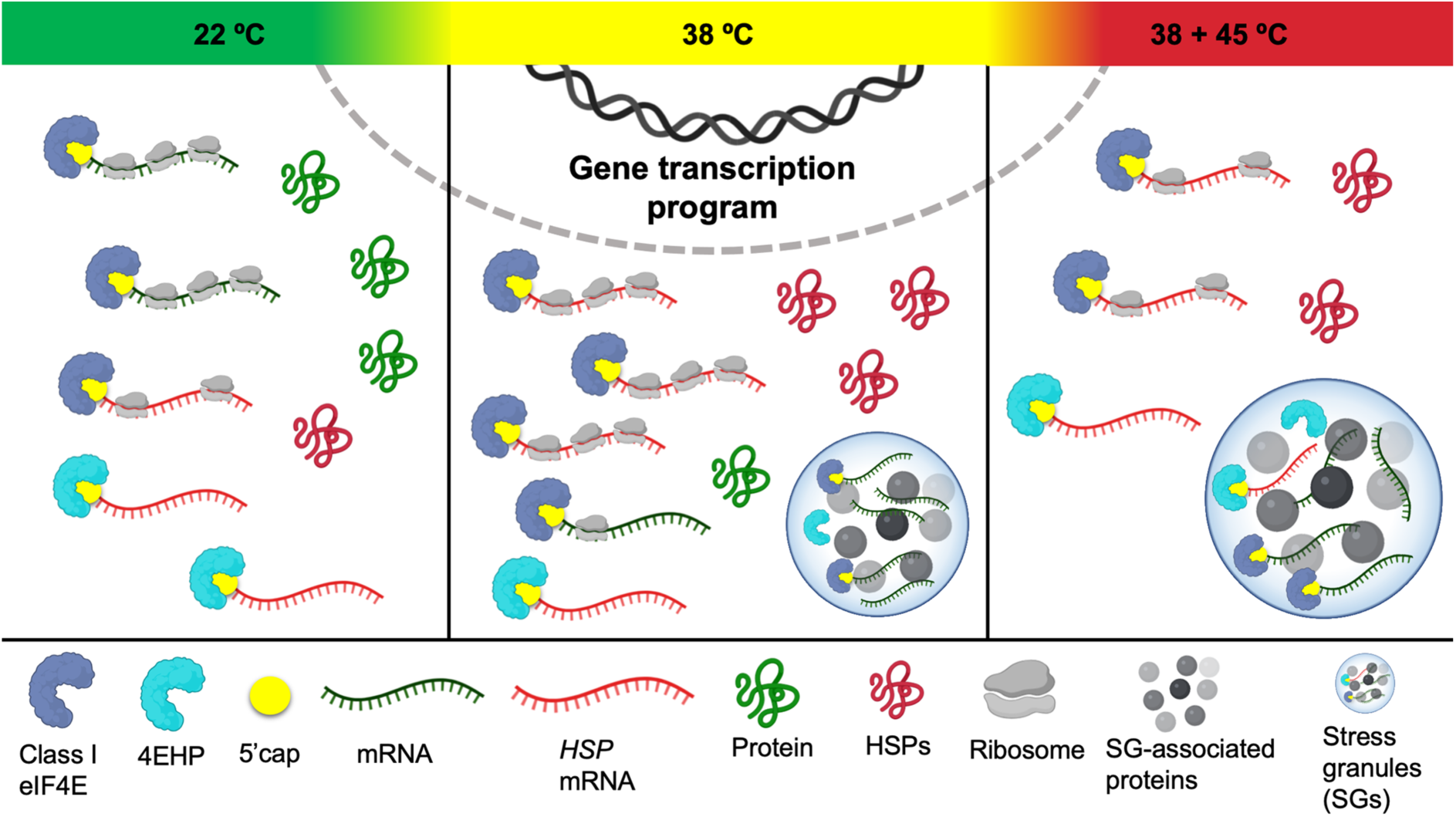
A proposed model for the role of 4EHP under control, acclimation, and heat stress (HS) conditions. At 22 °C, 4EHP is localized in the cytoplasm and modulates translation of basal *HSP* mRNAs, while Class I eIF4Es support general translation. During acclimation (38 °C), *HSP* transcripts accumulate and 4EHP becomes recruited to SGs to prevent their translation inhibition. A very high transcriptional activation of *HSP* mRNAs at this stage minimizes the inhibition by remaining cytoplasmic 4EHP. Upon HS of acclimated plants (38 + 45 °C), transcription of *HSP* is lowered and important translation inhibition takes place in the cytoplasm through increase of SG formation. At this point, cytoplasmic 4EHP binding to *HSP* mRNA leading to inhibition of its translation, SG sequestration and possibly degradation becomes visible, which is important to limit HSP abundance and facilitate restoration of growth as soon as temperature decreases.

In support of the model represented in **Fig. 8**, we found that the absence of 4EHP conduced to overaccumulation of more transcripts upon HS challenge, as compared to acclimation or normal temperatures. A subset of chaperone-encoding mRNAs remained at higher levels in *4ehp-1* than in Col-0 plants. Some of them displayed increased levels even under control conditions. Transcriptomic studies in plants have shown that HSP mRNA levels exhibit a biphasic pattern during prolonged heat exposure. They are rapidly induced at initial acclimation to moderate temperatures but decline when stress persists or during recovery, by activating mRNA decay pathways involving XRN4 and LARP1 ribonucleases (Charng *et al*., 2007; Merret *et al*., 2013; Bakery *et al*., 2024). In non-plant systems, HSP transcripts also show tightly controlled, transient induction rather than sustained overaccumulation in response to HS (Satyal *et al*., 1998; Tang *et al*., 2014; Quan *et al*., 2022). A controlled reduction of these transcripts prevents excessive HSP accumulation and allows cellular resource redirection to other protective pathways. The *4ehp-1* mutant presented elevated *HSP* transcript levels with respect to WT after severe HS. This indicates that 4EHP contributes to downregulation of HSP mRNAs during the HS phase. Since genes encoding heat-responsive transcription factors, which act as master regulators of the transcriptional response, were not significantly affected in the mutant line, deregulation of HSP mRNA levels is more likely a direct effect of the absence of 4EHP.

Translational efficiency was not significantly altered in the absence of 4EHP. The mutant displayed expected general inhibition of translation during heat shock, like WT plants. Four *HSP* mRNAs, *HSP101*, *HSP70B*, *HSP17.6A*, and *HSP17.6C* were preferentially localized to high and middle polyribosomal fractions under acclimation, but re-localized mainly to low polysomes and monosomes after HS in both, Col-0 and *4ehp-1*. It has been reported that translation elongation rates significantly vary among transcripts, and high ribosome occupancy does not always correlate with increased protein output, conferring a translational buffering that stabilizes protein levels despite changes in mRNA abundance (Zhao *et al*., 2019; Bai *et al*., 2021; Kusnadi *et al*., 2022). This mechanism is particularly relevant for HSPs, as highly inducible HSP70 mRNAs are enriched in non-optimal codons that promote ribosome stalling and activate ribosome quality control pathways under heat stress, thereby preventing deleterious HSP70 overaccumulation while still supporting the enhanced thermotolerance observed in the *4ehp-1* mutant (Alagar Boopathy *et al*., 2023). Post-translational mechanisms also buffer HSP accumulation, as ubiquitinated proteins generated during heat stress are rapidly targeted to the proteasome and proteasome activity remains elevated for many hours after heat shock (Maxwell *et al*., 2021; Lee & Goldberg, 2022). HSP101 itself interacts with the 26S proteasome and is dynamically recruited to cytosolic foci to promote aggregate disassembly (McLoughlin *et al*., 2019). Although essential for thermotolerance, HSP overaccumulation is known to produce growth and fitness costs, implying the existence of an optimal HSP abundance threshold (Queitsch *et al*., 2000; Tian *et al*., 2021).

According to the several layers of control, HSP protein levels are maintained within an optimal range throughout acclimation, short recovery, and heat stress. However, eventually posttranslational modifications lead to their decay and redistribution. Surprisingly, our shotgun proteomic analysis and the identification of differentially accumulated spots in 2D gels revealed a greater number and diversity of HSPs in *4ehp-1* in response to HS, but not under acclimation or normal temperature. Disturbance of HSP mRNA levels due to 4EHP absence might impact thermotolerance during the recovery period. On the other hand, the expression of a broader repertoire of HSPs at protein level (Supporting information Table S4) might contribute to the lower sensitivity to HS of the mutant. The effect of such a pattern is supported by previous studies highlighting the importance of chaperone diversity in thermotolerance response (Lin *et al*., 1984; Krishnan *et al*., 1989). For example, studies in transgenic rice (*Oryza sativa* cv. Hoshinoyume) overexpressing HSP17.7 demonstrated increased heat tolerance and UV resistance compared with non-transformed plants (Murakami *et al*., 2004). HSP17 proteins represent the largest proportion (42%) of HSPs in *A. thaliana* (Weston *et al*., 2011). We found significant increases for Class II HSP17.6A and Class I HSP17.6C at mRNA level, which was also identified in 2D-differential spots in *4ehp-1*. Hence, we propose that the enhanced HSP repertoire in the mutant contributes to its greater resistance to thermal stress.

Finally, the observation of differentially expressed genes associated with cellular responses to hypoxia in the *4ehp-1* mutant suggests that 4EHP also contributes to proper tolerance to this type of stress. It would be particularly relevant to examine its subcellular localization under low oxygen conditions and mutant tolerance to such stress. This could reveal additional roles of plant 4EHP aligning with some functions of its human ortholog (Ho *et al*., 2016).

In summary, this work provides new insights into the role of 4EHP at multiple levels of molecular regulation. Our findings support a regulatory model where cytoplasmic 4EHP likely acts as a direct repressor, binding target transcripts to inhibit their translation and stability. However, because it functions as a cap-binding protein, we propose that 4EHP could also actively recruit its interacting mRNAs into stress granules (SGs) in response to heat stress (Figure 8). This dynamic recruitment modulates the availability of heat stress-related mRNAs for translation, explaining why its absence is associated with a broader repertoire of heat-responsive proteins and enhanced plant thermotolerance. Previous studies describing the resistance of the 4EHP mutant to virus infection (Gomez *et al*., 2019; Suzuki *et al*., 2025) and improved resistance to *P. xylostella* larvae (Lin et al 2025) together with the enhanced thermotolerance reported here, make it a promising candidate for targeted genome editing for improved growth and stress tolerance in crop plants.

## Data availability

The RNA-seq data sets are deposited in the National Center for Biotechnology Information (NCBI) Gene Expression Omnibus (GEO) under the accession number GSE326555. All transcript and protein data generated or analyzed in this study are presented in the figures and tables of this manuscript and its Supporting Information files.

## Acknowledgements

We thank Dr. Víctor Zaldívar Machorro (USAII, Facultad de Química, UNAM) for his technical support with the shotgun proteomic analysis. We are also grateful for the technical assistance provided by Wiebke Hellmeyer and Judith Mehrmann (Institute of Plant Sciences and Microbiology, University of Hamburg), and M.Sc. Ana Laura Guzmán Ortiz (Unidad de Investigación en Inmunología y Proteómica, Hospital Infantil de México Federico Gómez).

## Author contributions

MAPT, KQN, MW and TDD conceptualized the research; MAPT, KQN, EDC, STD, ALD, MNM, OH and JHD developed the different methods and collected the data; MAPT, KQN, JHD, HQ and TDD curated and analyzed the data; MAPT, MW and TDD elaborated the figures. MAPT and TDD wrote the original draft of the manuscript. All co-authors revised and edited the manuscript.

## REFERENCES

Alagar Boopathy LR, Beadle E, Xiao AR, Garcia-Bueno Rico A, Alecki C, Garcia de-Andres I, Edelmeier K, Lazzari L, Amiri M, Vera M. 2023. The ribosome quality control factor Asc1 determines the fate of HSP70 mRNA on and off the ribosome. Nucleic Acids Res 51(12): 6370–6388.

Alonso JM, Stepanova AN, Leisse TJ, Kim CJ, Chen H, Shinn P, Stevenson DK, Zimmerman J, Barajas P, Cheuk R, et al. 2003. Genome-wide insertional mutagenesis of Arabidopsis thaliana. Science 301(5633): 653–657.

Bai B, van der Horst N, Cordewener JH, America AHP, Nijveen H, Bentsink L. 2021. Delayed Protein Changes During Seed Germination. Front Plant Sci 12: 735719.

Bakery A, Vraggalas S, Shalha B, Chauhan H, Benhamed M, Fragkostefanakis S. 2024. Heat stress transcription factors as the central molecular rheostat to optimize plant survival and recovery from heat stress. New Phytol 244(1): 51–64.

Bastet A, Zafirov D, Giovinazzo N, Guyon-Debast A, Nogue F, Robaglia C, Gallois JL. 2019. Mimicking natural polymorphism in eIF4E by CRISPR-Cas9 base editing is associated with resistance to potyviruses. Plant Biotechnol J 17(9): 1736–1750.

Browning KS, Bailey-Serres J. 2015. Mechanism of cytoplasmic mRNA translation. Arabidopsis Book 13: e0176.

Bush MS, Hutchins AP, Jones AM, Naldrett MJ, Jarmolowski A, Lloyd CW, Doonan JH. 2009. Selective recruitment of proteins to 5’ cap complexes during the growth cycle in Arabidopsis. Plant J 59(3): 400–412.

Chapat C, Jafarnejad SM, Matta-Camacho E, Hesketh GG, Gelbart IA, Attig J, Gkogkas CG, Alain T, Stern-Ginossar N, Fabian MR, et al. 2017. Cap-binding protein 4EHP effects translation silencing by microRNAs. Proc Natl Acad Sci U S A 114(21): 5425–5430.

Charng YY, Liu HC, Liu NY, Chi WT, Wang CN, Chang SH, Wang TT. 2007. A heat-inducible transcription factor, HsfA2, is required for extension of acquired thermotolerance in Arabidopsis. Plant Physiol 143(1): 251–262.

Chen B, Feder ME, Kang L. 2018. Evolution of heat-shock protein expression underlying adaptive responses to environmental stress. Mol Ecol 27(15): 3040–3054.

Chen H, Chen C, Huang S, Zhao M, Wang T, Jiang T, Wang C, Tao Z, Zhang Y, Wang Y, et al. 2023. Inactivation of RPX1 in Arabidopsis confers resistance to Plutella xylostella through the accumulation of the homoterpene DMNT. Plant Cell Environ 46(3): 946–961.

Chen R, Tu Z, He C, Nie X, Li K, Fei S, Song B, Nie B, Xie C. 2022. Susceptibility factor StEXA1 interacts with StnCBP to facilitate potato virus Y accumulation through the stress granule-dependent RNA regulatory pathway in potato. Horticulture Research.

Christie M, Igreja C. 2023. eIF4E-homologous protein (4EHP): a multifarious cap-binding protein. FEBS J 290(2): 266–285.

Dannfald A, Carpentier MC, Merret R, Favory JJ, Deragon JM. 2025. Plant response to intermittent heat stress involves modulation of mRNA translation efficiency. Plant Physiol 197(2).

Decker CJ, Parker R. 2012. P-bodies and stress granules: possible roles in the control of translation and mRNA degradation. Cold Spring Harb Perspect Biol 4(9): a012286.

Desroches Altamirano C, Kang MK, Jordan MA, Borianne T, Dilmen I, Gnadig M, von Appen A, Honigmann A, Franzmann TM, Alberti S. 2024. eIF4F is a thermo-sensing regulatory node in the translational heat shock response. Mol Cell 84(9): 1727–1741 e1712.

Ding Y, Shi Y, Yang S. 2020. Molecular Regulation of Plant Responses to Environmental Temperatures. Mol Plant 13(4): 544–564.

Doyle JJ, Doyle JL. 1987. A Rapid DNA Isolation Procedure for Small Quantities of Fresh Leaf Tissue. Phytochemical Bulletin 19: 11–15.

Duprat A, Caranta C, Revers F, Menand B, Browning KS, Robaglia C. 2002. The Arabidopsis eukaryotic initiation factor (iso)4E is dispensable for plant growth but required for susceptibility to potyviruses. Plant J 32(6): 927–934.

Earley KW, Haag JR, Pontes O, Opper K, Juehne T, Song K, Pikaard CS. 2006. Gateway-compatible vectors for plant functional genomics and proteomics. Plant J 45(4): 616–629.

Echevarria-Zomeno S, Yanguez E, Fernandez-Bautista N, Castro-Sanz AB, Ferrando A, Castellano MM. 2013. Regulation of Translation Initiation under Biotic and Abiotic Stresses. Int J Mol Sci 14(3): 4670–4683.

Frydryskova K, Masek T, Borcin K, Mrvova S, Venturi V, Pospisek M. 2016. Distinct recruitment of human eIF4E isoforms to processing bodies and stress granules. BMC Mol Biol 17(1): 21.

Galaxy C. 2024. The Galaxy platform for accessible, reproducible, and collaborative data analyses: 2024 update. Nucleic Acids Res 52(W1): W83–W94.

Ge SX, Jung D, Yao R. 2020. ShinyGO: a graphical gene-set enrichment tool for animals and plants. Bioinformatics 36(8): 2628–2629.

Gomez MA, Lin ZD, Moll T, Chauhan RD, Hayden L, Renninger K, Beyene G, Taylor NJ, Carrington JC, Staskawicz BJ, et al. 2019. Simultaneous CRISPR/Cas9-mediated editing of cassava eIF4E isoforms nCBP-1 and nCBP-2 reduces cassava brown streak disease symptom severity and incidence. Plant Biotechnol J 17(2): 421–434.

Gutierrez-Beltran E, Elander PH, Dalman K, Dayhoff GW, 2nd, Moschou PN, Uversky VN, Crespo JL, Bozhkov PV. 2021. Tudor staphylococcal nuclease is a docking platform for stress granule components and is essential for SnRK1 activation in Arabidopsis. EMBO J 40(17): e105043.

Gutierrez-Beltran E, Moschou PN, Smertenko AP, Bozhkov PV. 2015. Tudor staphylococcal nuclease links formation of stress granules and processing bodies with mRNA catabolism in Arabidopsis. Plant Cell 27(3): 926–943.

Herrera-Diaz J, Jelezova MK, Cruz-Garcia F, Dinkova TD. 2018. Protein Disulfide Isomerase (PDI1-1) differential expression and modification in Mexican malting barley cultivars. PLoS One 13(11): e0206470.

Ho JJD, Wang M, Audas TE, Kwon D, Carlsson SK, Timpano S, Evagelou SL, Brothers S, Gonzalgo ML, Krieger JR, et al. 2016. Systemic Reprogramming of Translation Efficiencies on Oxygen Stimulus. Cell Rep 14(6): 1293–1300.

Hong SW, Vierling E. 2000. Mutants of Arabidopsis thaliana defective in the acquisition of tolerance to high temperature stress. Proc Natl Acad Sci U S A 97(8): 4392–4397.

Hulsen T, de Vlieg J, Alkema W. 2008. BioVenn - a web application for the comparison and visualization of biological lists using area-proportional Venn diagrams. BMC Genomics 9: 488.

Joshi B, Lee K, Maeder DL, Jagus R. 2005. Phylogenetic analysis of eIF4E-family members. BMC Evol Biol 5: 48–68.

Kang Y, Lee K, Hoshikawa K, Kang M, Jang S. 2022. Molecular Bases of Heat Stress Responses in Vegetable Crops With Focusing on Heat Shock Factors and Heat Shock Proteins. Front Plant Sci 13: 837152.

Kazan K, Lyons R. 2016. The link between flowering time and stress tolerance. J Exp Bot 67(1): 47–60.

Keima T, Hagiwara-Komoda Y, Hashimoto M, Neriya Y, Koinuma H, Iwabuchi N, Nishida S, Yamaji Y, Namba S. 2017. Deficiency of the eIF4E isoform nCBP limits the cell-to-cell movement of a plant virus encoding triple-gene-block proteins in Arabidopsis thaliana. Sci Rep 7: 39678.

Kosmacz M, Gorka M, Schmidt S, Luzarowski M, Moreno JC, Szlachetko J, Leniak E, Sokolowska EM, Sofroni K, Schnittger A, et al. 2019. Protein and metabolite composition of Arabidopsis stress granules. New Phytol 222(3): 1420–1433.

Krishnan M, Nguyen HT, Burke JJ. 1989. Heat shock protein synthesis and thermal tolerance in wheat. Plant Physiol 90(1): 140–145.

Kropiwnicka A, Kuchta K, Lukaszewicz M, Kowalska J, Jemielity J, Ginalski K, Darzynkiewicz E, Zuberek J. 2015. Five eIF4E isoforms from Arabidopsis thaliana are characterized by distinct features of cap analogs binding. Biochem Biophys Res Commun 456(1): 47–52.

Kusnadi EP, Timpone C, Topisirovic I, Larsson O, Furic L. 2022. Regulation of gene expression via translational buffering. Biochim Biophys Acta Mol Cell Res 1869(1): 119140.

Lee D, Goldberg AL. 2022. 26S proteasomes become stably activated upon heat shock when ubiquitination and protein degradation increase. Proc Natl Acad Sci U S A 119(25): e2122482119.

Lin CY, Roberts JK, Key JL. 1984. Acquisition of Thermotolerance in Soybean Seedlings : Synthesis and Accumulation of Heat Shock Proteins and their Cellular Localization. Plant Physiol 74(1): 152–160.

Lin N, Ye H, Zhao M, Chen X, Ma J, Wang C, Wang T, Tao Z, Zhao Y, Zhang Q, et al. 2025. Glutathionylation-mediated degradation of a cap-binding protein enhances Arabidopsis resistance to Plutella xylostella. Plant Cell 37(8).

Lin Y, Protter DS, Rosen MK, Parker R. 2015. Formation and Maturation of Phase-Separated Liquid Droplets by RNA-Binding Proteins. Mol Cell 60(2): 208–219.

Livak KJ, Schmittgen TD. 2001. Analysis of relative gene expression data using real-time quantitative PCR and the 2(-Delta Delta C(T)) Method. Methods 25(4): 402–408.

Lohmann J, de Luxan-Hernandez C, Gao Y, Zoschke R, Weingartner M. 2023. Arabidopsis translation factor eEF1Bgamma impacts plant development and is associated with heat-induced cytoplasmic foci. J Exp Bot 74(8): 2585–2602.

Lohmann J, Herzog O, Rosenzweig K, Weingartner M. 2024. Thermal adaptation in plants: understanding the dynamics of translation factors and condensates. J Exp Bot 75(14): 4258–4273.

Marcotrigiano J, Gingras AC, Sonenberg N, Burley SK. 1997. Cocrystal structure of the messenger RNA 5’ cap-binding protein (eIF4E) bound to 7-methyl-GDP. Cell 89(6): 951–961.

Martinez-Silva AV, Aguirre-Martinez C, Flores-Tinoco CE, Alejandri-Ramirez ND, Dinkova TD. 2012. Translation initiation factor AteIF(iso)4E is involved in selective mRNA translation in Arabidopsis thaliana seedlings. PLoS One 7(2): e31606.

Maruri-Lopez I, Figueroa NE, Hernandez-Sanchez IE, Chodasiewicz M. 2021. Plant Stress Granules: Trends and Beyond. Front Plant Sci 12: 722643.

Matsuo H, Li H, McGuire AM, Fletcher CM, Gingras AC, Sonenberg N, Wagner G. 1997. Structure of translation factor eIF4E bound to m7GDP and interaction with 4E-binding protein. Nat Struct Biol 4(9): 717–724.

Maxwell BA, Gwon Y, Mishra A, Peng J, Nakamura H, Zhang K, Kim HJ, Taylor JP. 2021. Ubiquitination is essential for recovery of cellular activities after heat shock. Science 372(6549): eabc3593.

McLoughlin F, Kim M, Marshall RS, Vierstra RD, Vierling E. 2019. HSP101 Interacts with the Proteasome and Promotes the Clearance of Ubiquitylated Protein Aggregates. Plant Physiol 180(4): 1829–1847.

Merret R, Carpentier MC, Favory JJ, Picart C, Descombin J, Bousquet-Antonelli C, Tillard P, Lejay L, Deragon JM, Charng YY. 2017. Heat Shock Protein HSP101 Affects the Release of Ribosomal Protein mRNAs for Recovery after Heat Shock. Plant Physiol 174(2): 1216–1225.

Merret R, Descombin J, Juan YT, Favory JJ, Carpentier MC, Chaparro C, Charng YY, Deragon JM, Bousquet-Antonelli C. 2013. XRN4 and LARP1 are required for a heat-triggered mRNA decay pathway involved in plant acclimation and survival during thermal stress. Cell Rep 5(5): 1279–1293.

Morita M, Ler LW, Fabian MR, Siddiqui N, Mullin M, Henderson VC, Alain T, Fonseca BD, Karashchuk G, Bennett CF, et al. 2012. A novel 4EHP-GIGYF2 translational repressor complex is essential for mammalian development. Mol Cell Biol 32(17): 3585–3593.

Murakami T, Matsuba S, Funatsuki H, Kawaguchi K, Saruyama H, Tanida M, Sato Y. 2004. Over-expression of a small heat shock protein, sHSP17. 7, confers both heat tolerance and UV-B resistance to rice plants. Molecular Breeding 13(2): 165–175.

Nishikawa M, Katsu K, Koinuma H, Hashimoto M, Neriya Y, Matsuyama J, Yamamoto T, Suzuki M, Matsumoto O, Matsui H, et al. 2023. Interaction of EXA1 and eIF4E Family Members Facilitates Potexvirus Infection in Arabidopsis thaliana. J Virol 97(6): e0022123.

Ohama N, Kusakabe K, Mizoi J, Zhao H, Kidokoro S, Koizumi S, Takahashi F, Ishida T, Yanagisawa S, Shinozaki K, et al. 2016. The Transcriptional Cascade in the Heat Stress Response of Arabidopsis Is Strictly Regulated at the Level of Transcription Factor Expression. Plant Cell 28(1): 181–201.

Preiss T, M WH. 2003. Starting the protein synthesis machine: eukaryotic translation initiation. Bioessays 25(12): 1201–1211.

Preston JC, Fjellheim S. 2022. Flowering time runs hot and cold. Plant Physiol 190(1): 5–18.

Protter DSW, Rao BS, Van Treeck B, Lin Y, Mizoue L, Rosen MK, Parker R. 2018. Intrinsically Disordered Regions Can Contribute Promiscuous Interactions to RNP Granule Assembly. Cell Rep 22(6): 1401–1412.

Quan Y, Wang Z, Wei H, He K. 2022. Transcription dynamics of heat shock proteins in response to thermal acclimation in Ostrinia furnacalis. Front Physiol 13: 992293.

Queitsch C, Hong SW, Vierling E, Lindquist S. 2000. Heat shock protein 101 plays a crucial role in thermotolerance in Arabidopsis. Plant Cell 12(4): 479–492.

Rao KVM 2006. Introduction. In: Rao KVM, Raghavendra AS, Reddy KJ eds. Physiology and molecular biology of stress tolerance. Dordrecht, Netherlands: Springer, 1–14.

Rosettani P, Knapp S, Vismara MG, Rusconi L, Cameron AD. 2007. Structures of the human eIF4E homologous protein, h4EHP, in its m7GTP-bound and unliganded forms. J Mol Biol 368(3): 691–705.

Ruud KA, Kuhlow C, Goss DJ, Browning KS. 1998. Identification and characterization of a novel cap-binding protein from Arabidopsis thaliana. J Biol Chem 273(17): 10325–10330.

Salazar-Diaz K, Aquino-Luna M, Hernandez-Lucero E, Nieto-Rivera B, Pulido-Torres MA, Jorge-Perez JH, Gavilanes-Ruiz M, Dinkova TD. 2021. Arabidopsis thaliana eIF4E1 and eIF(iso)4E Participate in Cold Response and Promote Translation of Some Stress-Related mRNAs. Front Plant Sci 12: 698585.

Satyal SH, Chen D, Fox SG, Kramer JM, Morimoto RI. 1998. Negative regulation of the heat shock transcriptional response by HSBP1. Genes Dev 12(13): 1962–1974.

Scheer H, de Almeida C, Ferrier E, Simonnot Q, Poirier L, Pflieger D, Sement FM, Koechler S, Piermaria C, Krawczyk P, et al. 2021. The TUTase URT1 connects decapping activators and prevents the accumulation of excessively deadenylated mRNAs to avoid siRNA biogenesis. Nat Commun 12(1): 1298.

Schneider CA, Rasband WS, Eliceiri KW. 2012. NIH Image to ImageJ: 25 years of image analysis. Nat Methods 9(7): 671–675.

Schwanhausser B, Busse D, Li N, Dittmar G, Schuchhardt J, Wolf J, Chen W, Selbach M. 2011. Global quantification of mammalian gene expression control. Nature 473(7347): 337–342.

Song H, Zhao R, Fan P, Wang X, Chen X, Li Y. 2009. Overexpression of AtHsp90.2, AtHsp90.5 and AtHsp90.7 in Arabidopsis thaliana enhances plant sensitivity to salt and drought stresses. Planta 229(4): 955–964.

Staacke T, Mueller-Roeber B, Balazadeh S. 2025. Stress resilience in plants: the complex interplay between heat stress memory and resetting. New Phytol 245(6): 2402–2421.

Sun X, Sun C, Li Z, Hu Q, Han L, Luo H. 2016. AsHSP17, a creeping bentgrass small heat shock protein modulates plant photosynthesis and ABA-dependent and independent signalling to attenuate plant response to abiotic stress. Plant Cell Environ 39(6): 1320–1337.

Suzuki M, Nishikawa M, Yamamoto T, Koinuma H, Keima T, Fujimoto Y, Komatsu K, Hashimoto M, Neriya Y, Maejima K, et al. 2025. Broadening Virus Resistance Through Gene Pyramiding of eIF4E Family Members. Mol Plant Pathol 26(12): e70187.

Tang S, Lv Y, Chen H, Adam A, Cheng Y, Hartung J, Bao E. 2014. Comparative analysis of alphaB-crystallin expression in heat-stressed myocardial cells in vivo and in vitro. PLoS One 9(1): e86937.

Tian F, Hu XL, Yao T, Yang X, Chen JG, Lu MZ, Zhang J. 2021. Recent Advances in the Roles of HSFs and HSPs in Heat Stress Response in Woody Plants. Front Plant Sci 12: 704905.

Villaescusa JC, Buratti C, Penkov D, Mathiasen L, Planaguma J, Ferretti E, Blasi F. 2009. Cytoplasmic Prep1 interacts with 4EHP inhibiting Hoxb4 translation. PLoS One 4(4): e5213.

Wang TY, Liu JF, Chu ZY, Zhao YB, Ma J, Tao Z, Wang CH, Liu LZ, Li PJ. 2025. A genome-wide association study uncovers that BnaA10.NCBP regulates early flowering in. Industrial Crops and Products 226.

Weber C, Nover L, Fauth M. 2008. Plant stress granules and mRNA processing bodies are distinct from heat stress granules. Plant J 56(4): 517–530.

Weston DJ, Karve AA, Gunter LE, Jawdy SS, Yang X, Allen SM, Wullschleger SD. 2011. Comparative physiology and transcriptional networks underlying the heat shock response in Populus trichocarpa, Arabidopsis thaliana and Glycine max. Plant Cell Environ 34(9): 1488–1506.

Wu HL, Jen J, Hsu PY. 2024. What, where, and how: Regulation of translation and the translational landscape in plants. Plant Cell 36(5): 1540–1564.

Yanguez E, Castro-Sanz AB, Fernandez-Bautista N, Oliveros JC, Castellano MM. 2013. Analysis of genome-wide changes in the translatome of Arabidopsis seedlings subjected to heat stress. PLoS One 8(8): e71425.

Yoo SD, Cho YH, Sheen J. 2007. Arabidopsis mesophyll protoplasts: a versatile cell system for transient gene expression analysis. Nat Protoc 2(7): 1565–1572.

Zhang J, Jin H, Chen Y, Jiang Y, Gu L, Lin G, Lin C, Wang Q. 2024. The eukaryotic translation initiation factor eIF4E regulates flowering and circadian rhythm in Arabidopsis. Plant J 120(1): 123–138.

Zhao J, Qin B, Nikolay R, Spahn CMT, Zhang G. 2019. Translatomics: The Global View of Translation. Int J Mol Sci 20(1).

