## Supporting Information for "The eIF4E-homologous protein 4EHP (nCBP) regulates thermotolerance by modulating heat-responsive mRNAs and the HSP repertoire in Arabidopsis thaliana"

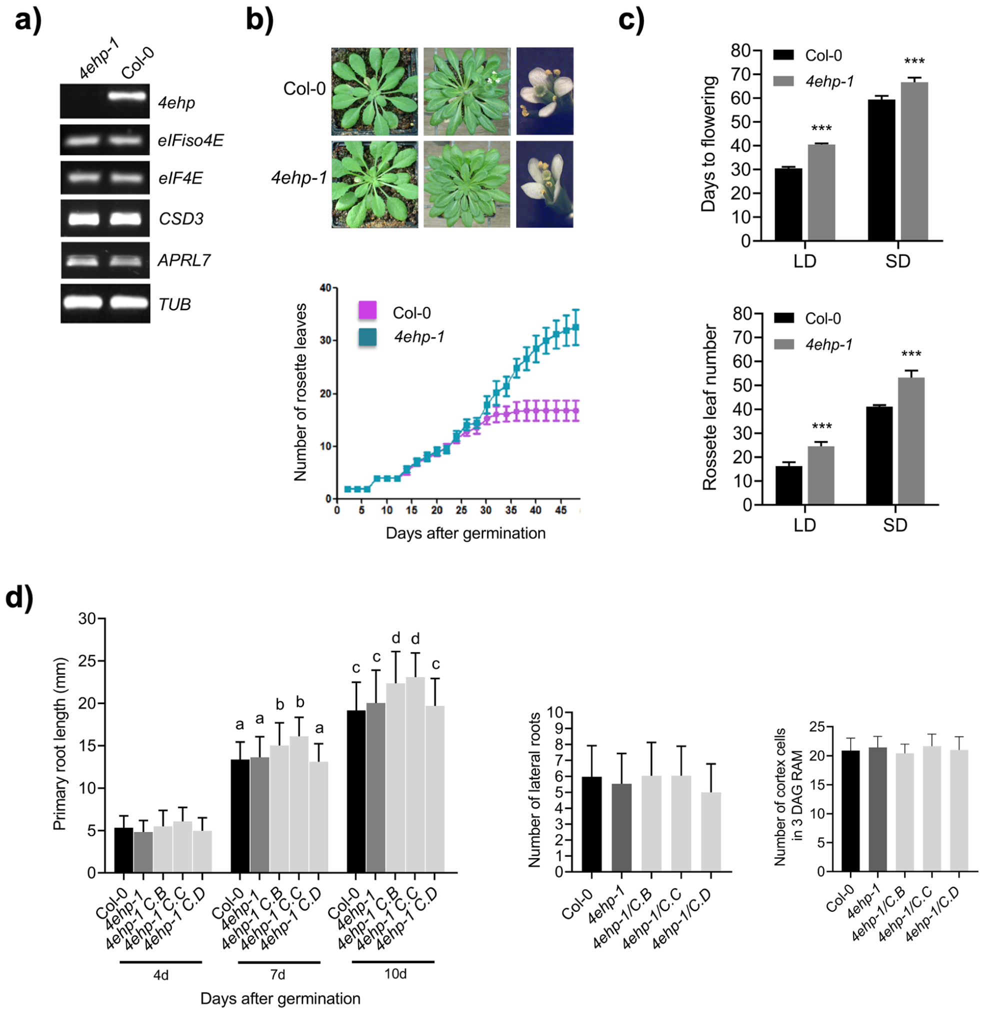


**Figure S1. Additional characterization of the *4ehp-1* mutant line.**
**a)** Reverse transcription – end point PCR analysis of the *4ehp-1* mutant using total RNA from 15-days-old seedlings. The *4EHP* transcript was not detected in the mutant, while the levels of *eIF4E* and *eIF(iso)4E* transcripts or those corresponding to genes bordering the chromosomal location of *4EHP* (*APRL7* and *CSD3*) were not affected. Alpha-Tubulin (*TUB*) was used as a loading control.
**b)** and **c)** Flowering Phenotype of wild-type (Col-0) and mutant (*4ehp-1*) plants grown under short day (SD) or long day (LD) conditions. Images correspond to overall morphology during vegetative growth and flowering. The number of rosette leaves was registered upon germination until bolting in three independent experiments with n=30. Bars represent standard error; asterisks indicate statistically significant difference at P<0.001.
**d)** Root growth was analyzed at the indicated time points using ImageJ software. Quantification of lateral root (including primordia) was performed on 10-days-old seedlings. Quantification of dividing cells in the cortical cell layer of the root apical meristem (RAM) was performed on 3-days-old seedlings. For the left and central panels, data represent mean ± SD from 60-90 roots per genotype divided into five biological replicates. Different letters indicate statistically significant differences (one-way ANOVA with Tukey's post-hoc test, P < 0.05). For the right panel, data represent mean ± SD from 20-30 roots per genotype divided into four biological replicates.

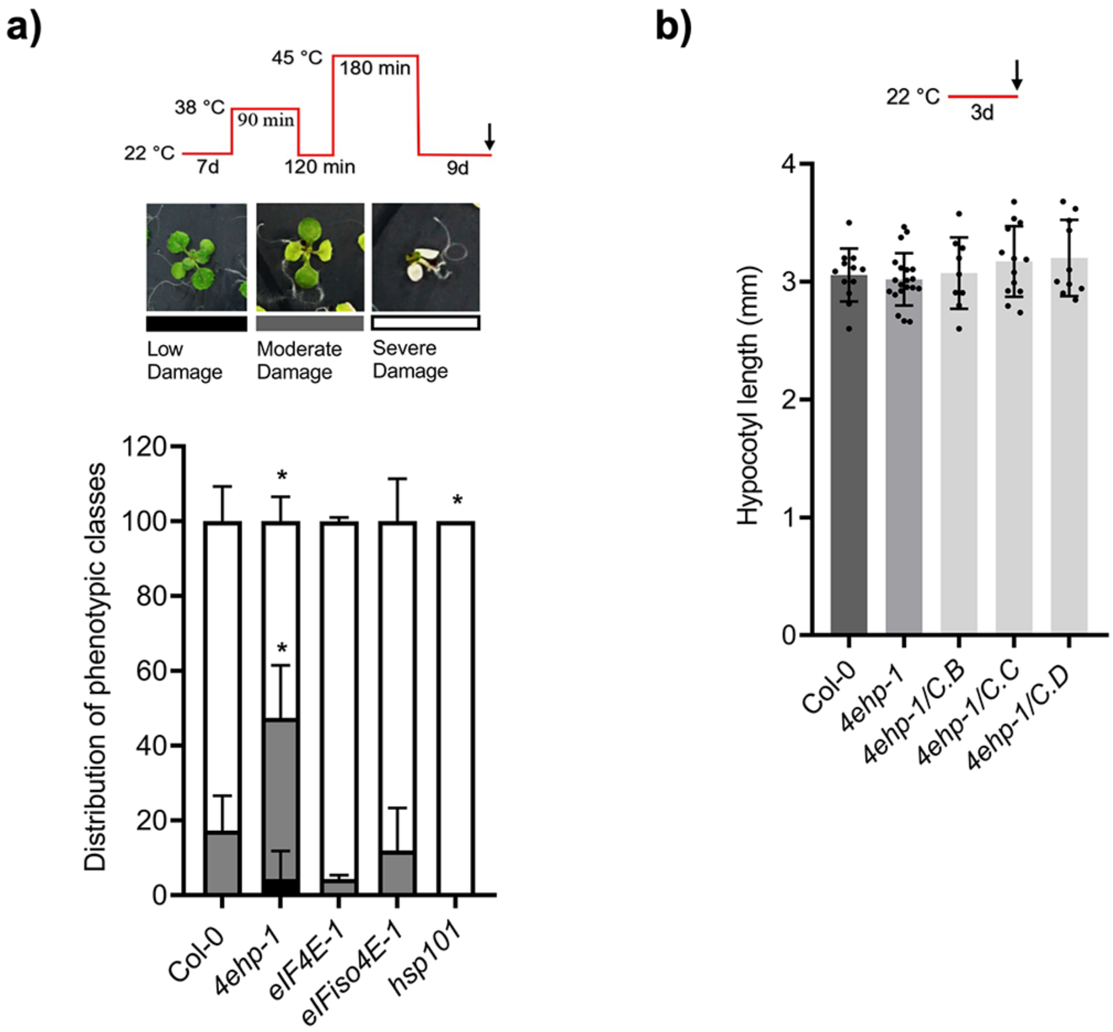


**Figure S2. Thermotolerance comparison between *eif4e* mutant lines and Col-0.**
**a)** Acquired thermotolerance was evaluated after 9 days of recovery from heat stress (45ºC) applied to 7-days-old seedlings previously acclimated at 38ºC. Three phenotypes were registered: low (black bars), moderate (grey bars), or severe (white bars) damage, according to the number of expanded green leaves, including cotyledons. Data represent mean ± SD from 100–160 seedlings per genotype divided into three biological replicates. * Significant differences compared with Col-0 (two-tailed Student's t-test; $P < 0.05$).
**b)** Hypocotyl elongation of 3-day-old seedlings grown under dark conditions at 22ºC. No differences were observed between Col-0, the *4ehp-1* mutant and three complemented lines (C.B, C.C, and C.D).

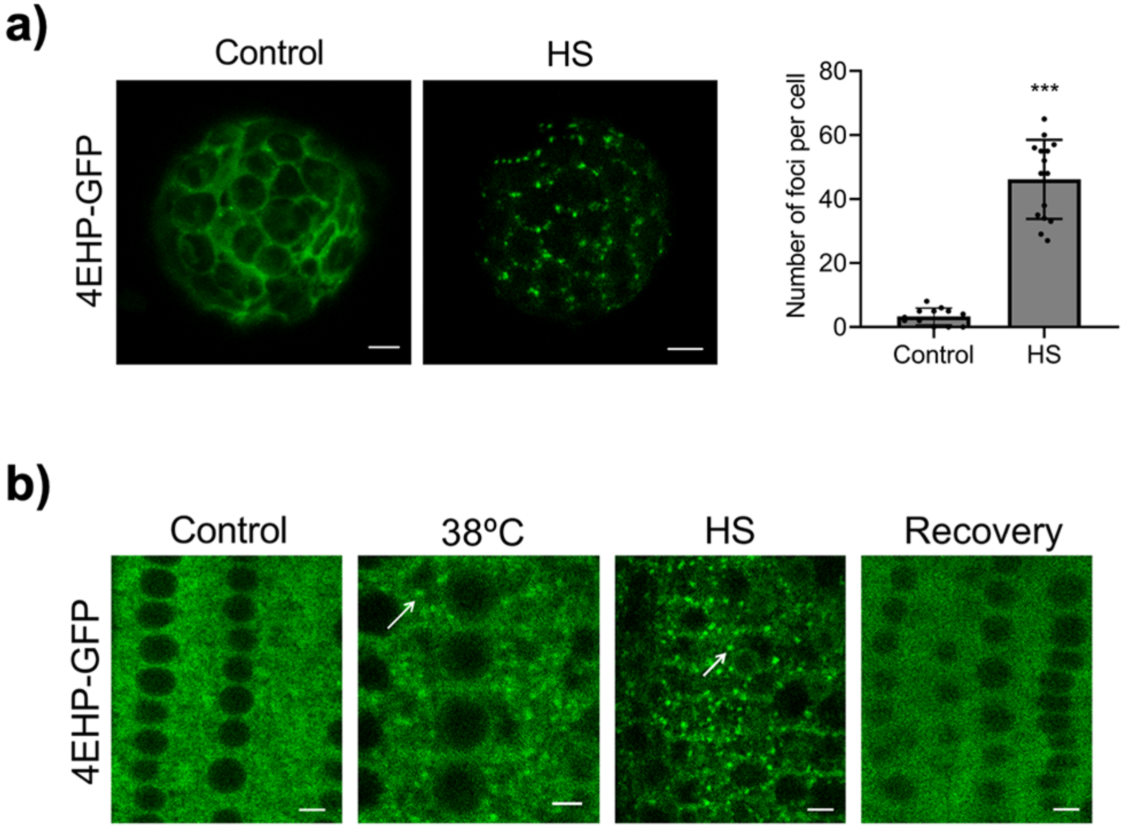


**Figure S3. Recruitment of 4EHP to stress granules during HS using a construct under its own promoter.**
**a)** Confocal images of protoplasts transiently expressing p4EHP:4EHP-GFP under control conditions or after heat treatment at 40°C for 35 min (HS). The right graph represents quantification of foci number per cell in control and HS protoplasts.
*** Significant difference between treatments n=15 (two-tailed Student's t-test; $P < 0.001$). Scale bar = 5 µm.
**b)** Localization of 4EHP-GFP in root cells of five-days-old plants of a mutant line complemented with p4EHP:4EHP-GFP. Seedlings were grown at 22°C as control or incubated at 38ºC for 1.5h or at 40°C for 35 min (HS). The recovery period was for 6 h, at control temperature after HS. White arrows indicate 4EHP-GFP accumulation in cytoplasmic foci. Scale bar = 5 µm.

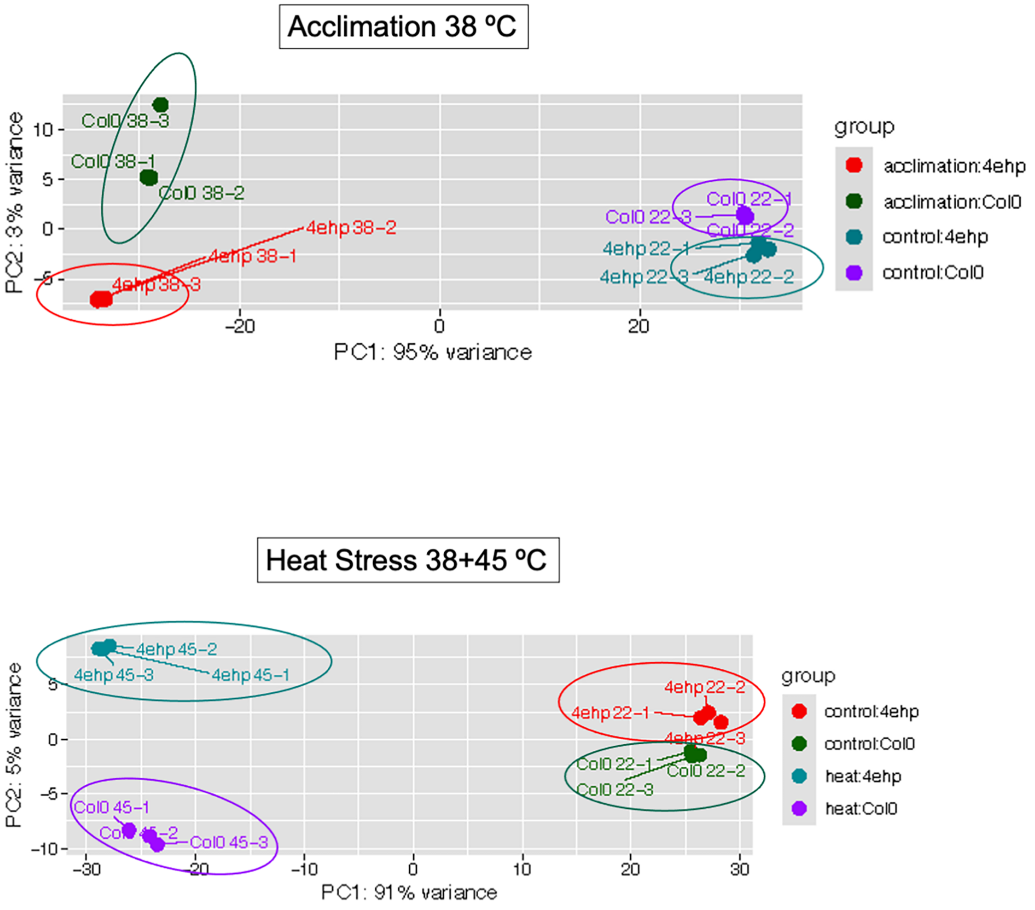


**Figure S4. Principal component analysis (PCA) for transcriptomes from *4ehp-1* and Col-0 at different temperatures.**
PC1 scores difference between temperatures, while PC2 scores difference between lines. Data for the control temperature are shown in both panels. Three biological replicates are circled for each condition and line.

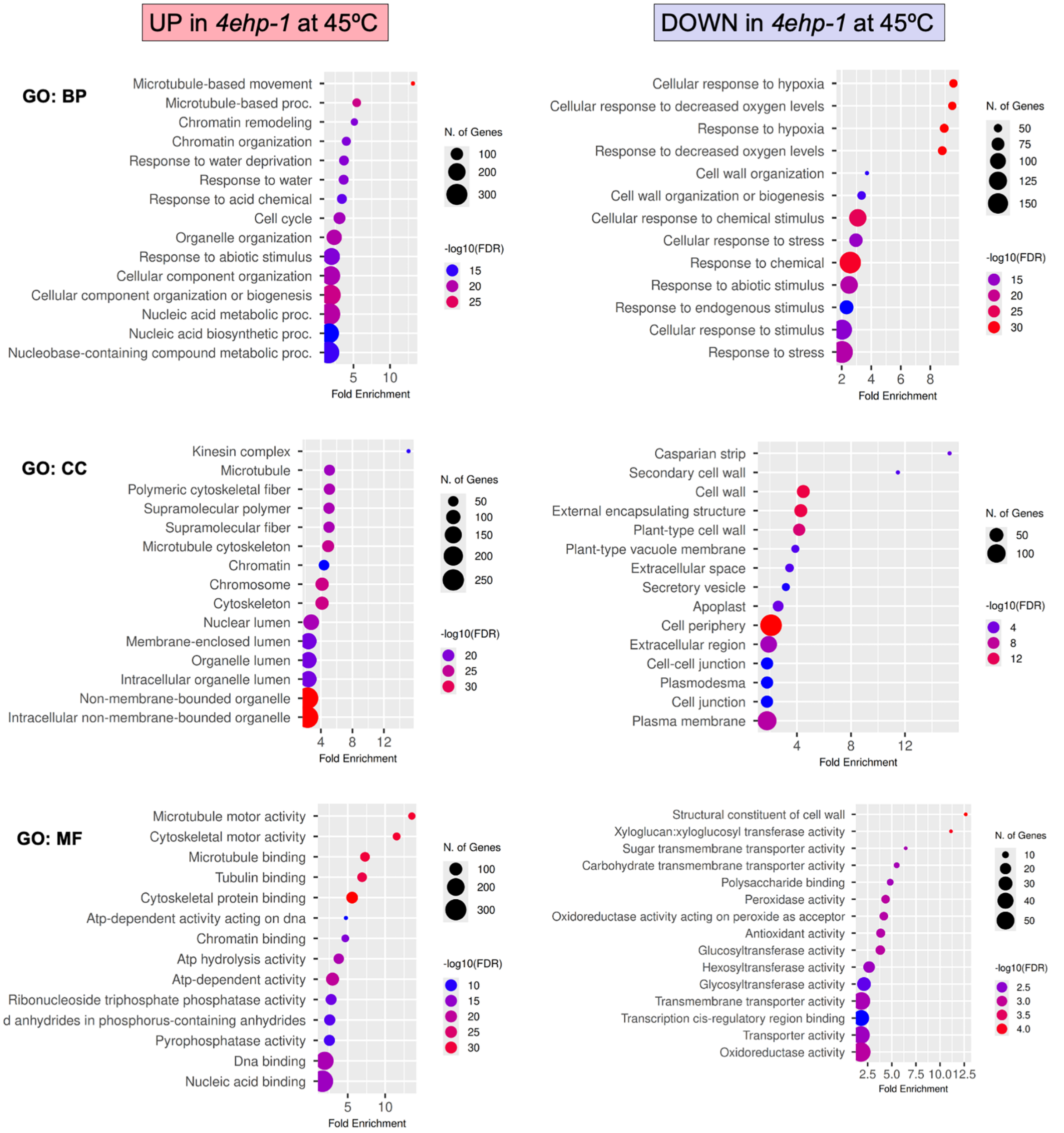


**Figure S5. GO enrichment analysis of upregulated and down-regulated transcripts in *4ehp-1* upon HS.**

The most significantly enriched GO terms with -log10(FDR) < 5 are presented for biological process (BP), cellular component (CC), and molecular function (MF). The interpretation of dot sizes and colors is shown at left for each plot. Plots were generated in ShinyGO 0.85.

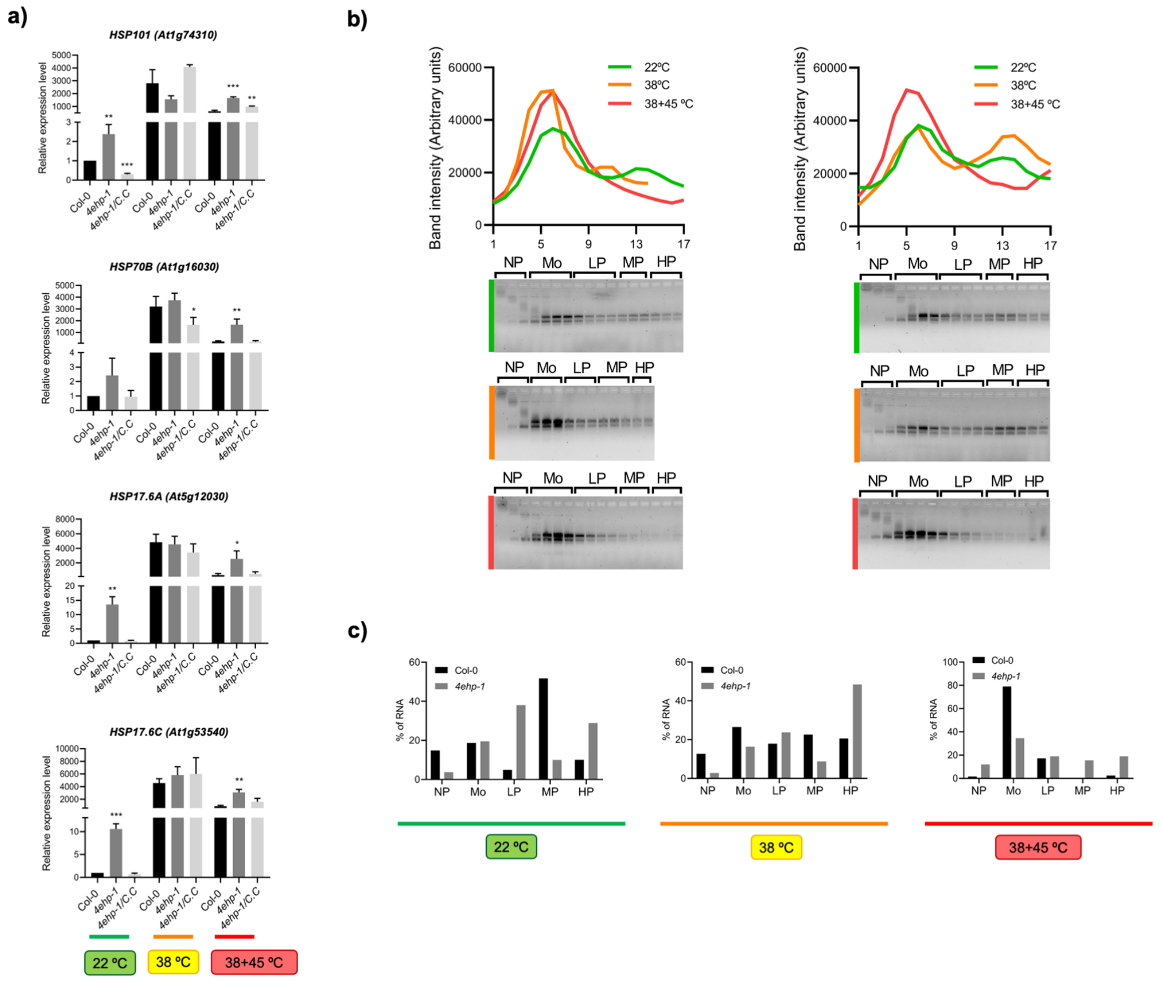


**Figure S6.** ***HSP* transcript levels in complemented *4ehp-1* lines and polyribosomal profiling.**

**a)** The accumulation of *HSP101*, *HSP70B*, *HSP17.6A* and *HSP17.6C* is shown for the three different temperatures (control, 22 ºC; acclimation, 38 ºC; heat stress after recovery, 38+45 ºC) in Col-0, *4ehp-1* and complemented line C seedlings by RT-qPCR. *Actin2* was used for normalization and Col-0 at 22 ºC as reference. Results are mean of three independent biological replicates. Significance of differential accumulation is denoted by: (*) $p<0.05$; (**) $p<0.01$; (***) $p<0.001$.

**b)** Col-0 and *4ehp-1* polyribosomal profiles at each temperature showing all fractions collected from the sucrose density gradient analyzed by gel electrophoresis. The fractions pooled for transcript analysis are indicated: non-polysome (NP), monosome (M), low polysome (LP), mid polysome (MP) and high polysome (HP).

**c)** Distribution of *Actin2* along the pooled polyribosomal fractions at each temperature for Col-0 and *4ehp-1* seedlings.

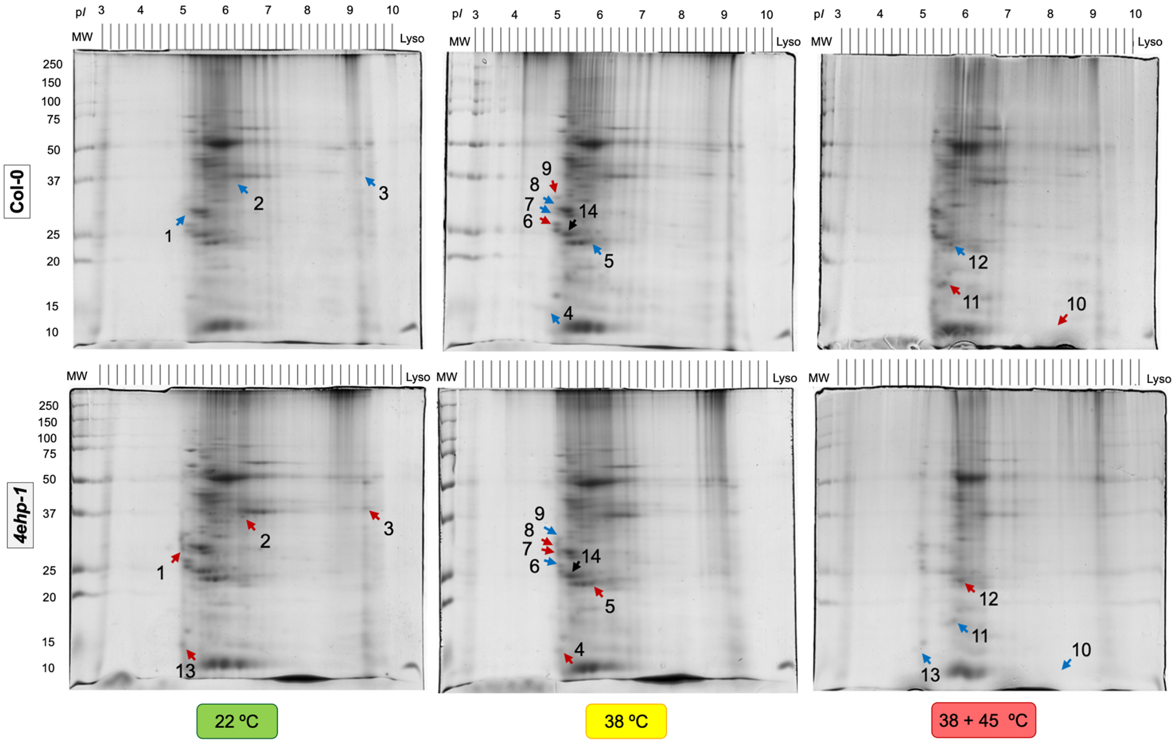


**Figure S7. Proteomic profiles by two-dimensional gel maps.**
Proteins were extracted from Col-0 and *4ehp-1* mutant seedlings grown for 7 days at 22 °C under long day photoperiod (control, 22 °C), subjected to 38 °C for 90 min (acclimation, 38 °C), recovered for 2 h at 22 °C and then incubated at 45 °C for 2 h (HS, 38 + 45 °C). Equal amount (350 μg) of total proteins were separated on IPG strips pH 3−10 (first dimension) and then on 12% SDS-PAGE (second dimension). The gels were stained with colloidal Coomassie Blue G-250 and images were acquired on a calibrated densitometer (BioRad). Differential protein spots between Col-0 and *4ehp-1* at each temperature are highlighted by arrows (red, increased; blue, decreased). A spot with similar intensity (spot 14) is indicated by black arrow. Spots were numbered for identification by LC/MS/MS on each gel. MW, molecular weight marker with masses in kDa shown at left for each gel; Lyso, 1 μg of lysozyme was used as standard at the right lane of each gel.
